# Deep learning enables accurate clustering and batch effect removal in single-cell RNA-seq analysis

**DOI:** 10.1101/530378

**Authors:** Xiangjie Li, Yafei Lyu, Jihwan Park, Jingxiao Zhang, Dwight Stambolian, Katalin Susztak, Gang Hu, Mingyao Li

## Abstract

Single-cell RNA sequencing (scRNA-seq) can characterize cell types and states through unsupervised clustering, but the ever increasing number of cells imposes computational challenges. We present an unsupervised deep embedding algorithm for single-cell clustering (DESC) that iteratively learns cluster-specific gene expression signatures and cluster assignment. DESC significantly improves clustering accuracy across various datasets and is capable of removing complex batch effects while maintaining true biological variations.

---

A primary challenge in scRNA-seq analysis is analyzing the ever increasing number of cells, which can be thousands to millions in large projects such as the Human Cell Atlas^1^. Identifying cell populations is a challenge in large datasets because many existing scRNA-seq clustering methods cannot be scaled up to handle them. It is desirable to first learn cluster-specific gene expression features from cells that are easy to cluster because they provide valuable information on cluster-specific gene expression signatures. These cells can help improve clustering of cells that are hard-to-cluster.

Another challenge in scRNA-seq analysis is batch effect, which is systematic gene expression difference from one batch to another^2^. Batch effect is inevitable in studies involving human tissues because the data are often generated at different times and the batches can confound biological variations. Failure to remove batch effect will complicate downstream analysis and leads to a false interpretation of results.

ScRNA-seq clustering and batch effect removal are typically addressed through separate analyses. Commonly used approaches to remove batch effect include Seurat’s Canonical Correlation Analysis^3^ (CCA) or Mutual Nearest Neighbors (MNN) approach^4^. After removing batch effect, clustering analysis is performed to identify cell clusters using methods such as Louvain’s method^5^, Infomap^6^, graph-based clustering^7^, shared nearest neighbor^8^, or consensus clustering with SC3^9^. Since some cell types are more vulnerable to batch effect than others, batch effect removal should be performed jointly with clustering to achieve optimal performance. However, none of the existing methods are capable of simultaneously clustering cells and removing batch effect.

We developed DESC, an unsupervised deep learning algorithm that iteratively learns cluster-specific gene expression representation and cluster assignments for scRNA-seq data clustering (**Fig. 1a**). Using a deep neural network, DESC initializes clustering obtained from an autoencoder and learns a non-linear mapping function from the original scRNA-seq data space to a low-dimensional feature space by iteratively optimizing a clustering objective function. This iterative procedure moves each cell to its nearest cluster, balances biological and technical differences between clusters, and reduces the influence of batch effect. DESC also enables soft clustering by assigning cluster-specific probabilities to each cell, facilitating the clustering of cells with high-confidence.

**Figure 1.**
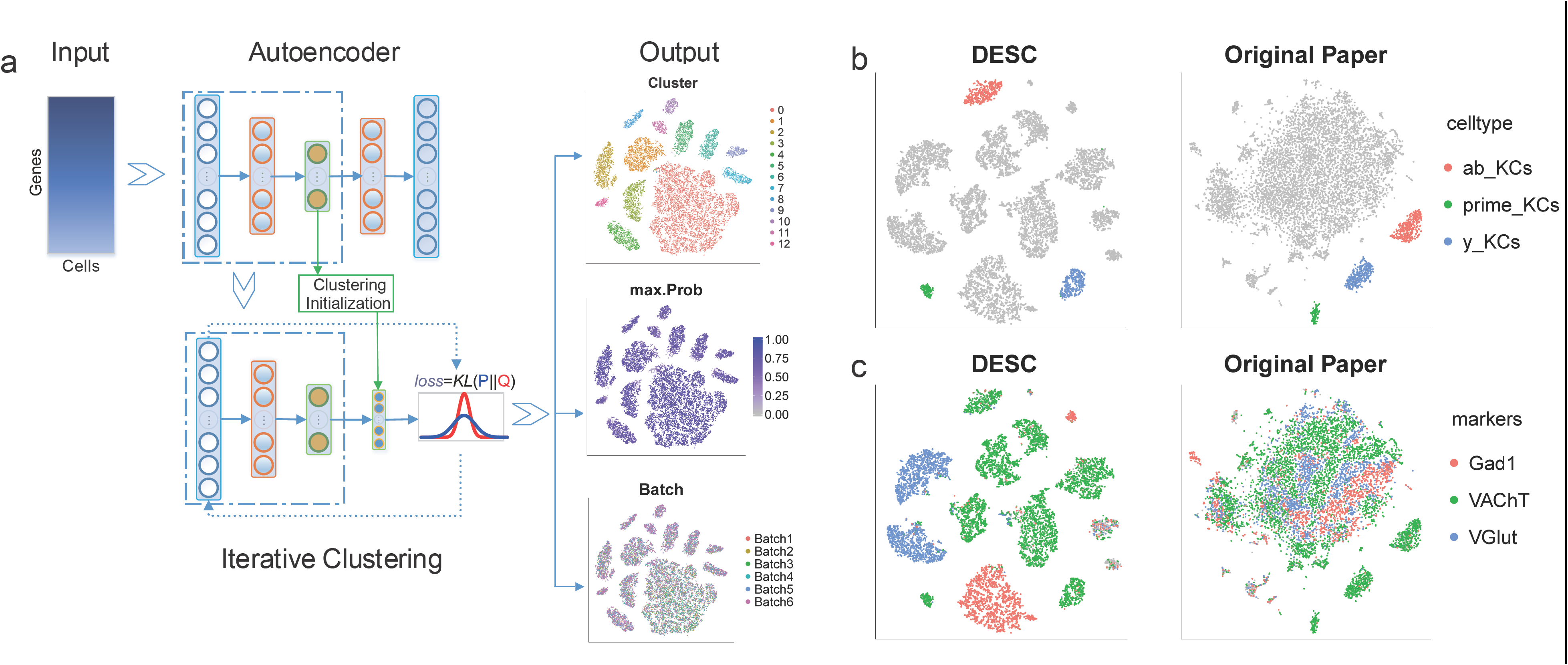
(a) Overview of the DESC framework. DESC starts with parameter initialization in which a stacked autoencoder is used for pretraining and learning a low-dimensional representation of the input gene expression matrix. The resulting encoder is then added to the iterative clustering neural network to cluster cells iteratively. The final output of DESC includes cluster assignment, the corresponding probabilities for cluster assignment for each cell, and the low-dimensional representation of the data. (b) Analysis of the single-cell data generated from midbrain in Drosophila. DESC not only identified the three types of Kenyon cells, which are detectable by the Louvain’s method, but also identified cholinergic, glutamatergic, and GABAergic neurons, which are harder-to-separate by the Louvain’s method reported in the original paper.

We benchmarked DESC’s performance by analyzing the multi-tissue gene expression data in GTEx^10^. We treat this dataset as the gold-standard because the tissue origins are known. Although not generated by scRNA-seq, GTEx data are similar to scRNA-seq in that it contains a large number of samples (n=11,688) originated from many tissue types and is similar to the volume and complexity of scRNA-seq data (**Supplementary Note 1**). DESC’s clustering yields an adjusted rand index (ARI) of 0.790, whereas the ARIs for Louvain’s method, SC3, and Infomap are 0.755, 0.349, and 0.267, respectively. As shown in the Sankey diagrams (**Supplementary Fig. 3**), samples that were misclassified by DESC and Louvain’s method tend to be from closely related tissues, whereas SC3 tends to misclassify samples from tissues distantly related.

We analyzed a scRNA-seq dataset generated from the midbrain of Drosophila, which includes 10,286 cells using Drop-seq^11^(**Supplementary Note 2**). This dataset has minimal batch effect. DESC identified three types of mushroom body Kenyons, with 1,038 out of the 1,053 Kenyon cells correctly classified, a 98.6% classification accuracy (**Fig. 1b**). DESC also separated cholinergic, glutamatergic, and GABAergic neurons, which were mixed together in the Louvain’s clustering as shown in the original paper (**Fig. 1c**). These results indicate that DESC can identify cell types that are detectable by the Louvain’s method, and is also able to separate more closely related cells, indicating its increased accuracy in classifying closely related cell types. We further applied Louvain’s method to the low-dimensional representation learned from the autoencoder in DESC for clustering, and separated the cholinergic, glutamatergic, and GABAergic neurons better than the original Louvain’s clustering with principal components (PC) based dimension reduction (**Supplementary Fig. 4**). These results suggest the autoencoder is more effective than PC in dimension reduction for single-cell clustering.

Encouraged by these findings, we analyzed a scRNA-seq dataset with known batch effect (**Supplementary Note 3**). Shekhar et al.^12^ sequenced 23,494 retinal bipolar cells using Drop-seq, where cells from six replicates were processed in two different batches. **Fig. 2a and 2c** show that DESC removed the batch effect, and yields an ARI of 0.973 for clustering. The corresponding ARIs for Louvain, SC3, Infomap, CCA and MNN are 0.965, 0.521, 0.560, 0.637, and 0.974, respectively. Although DESC, Louvain, and MNN have similar ARIs, DESC has the smallest Kullback-Leibler (KL) divergence, which measures the degree of random mixing of cells in different batches, indicating that DESC is more effective in removing batch effect (**Fig. 2b**). Further analysis revealed that the batch effect removal in DESC is due to its iterative clustering, in which cells from the same cluster, separated by technical batch effect, are grouped closer and closer to the cluster centroid over iterations (**Figs. 2d and 2e**).

**Figure 2.**
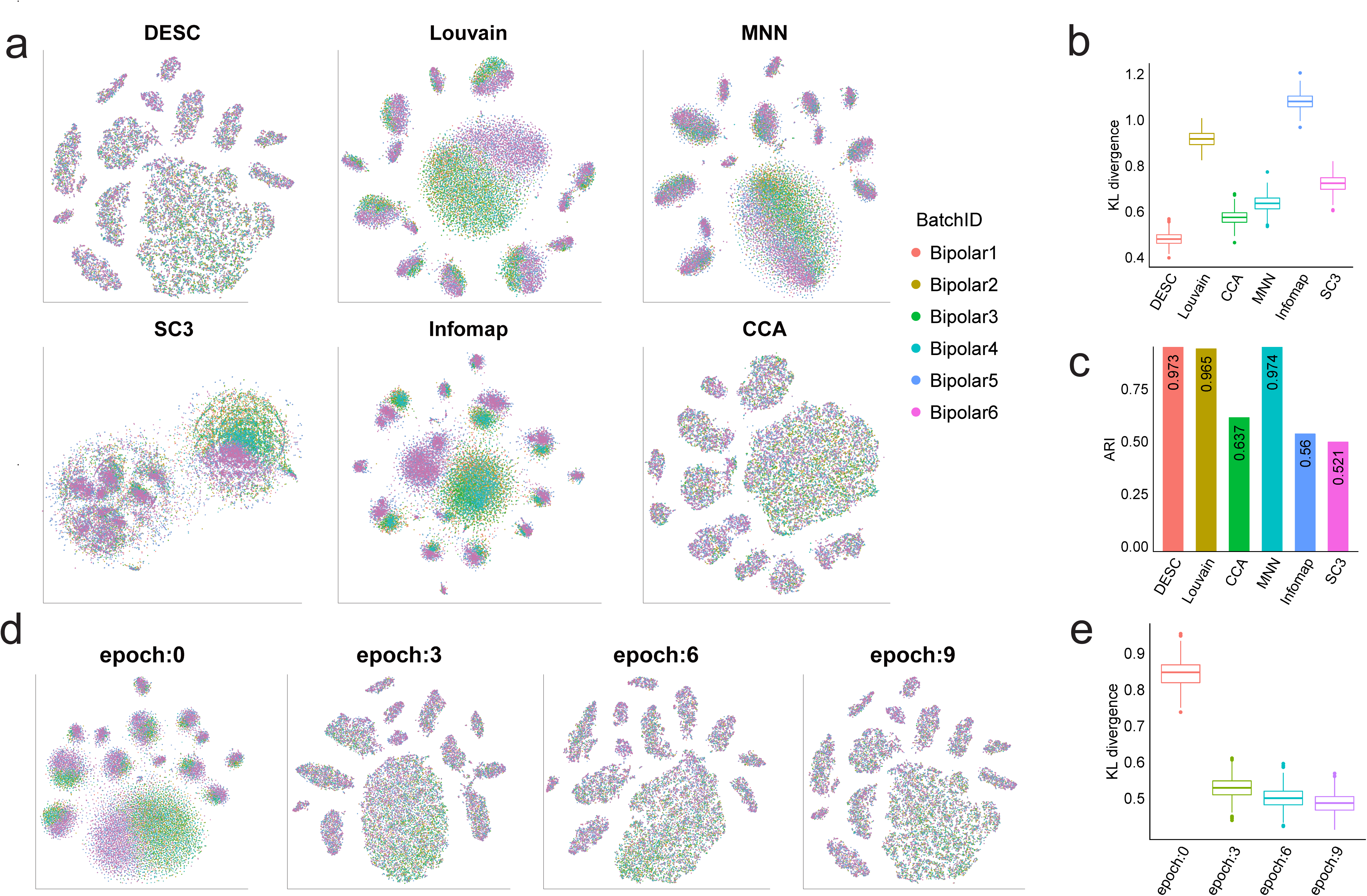
(a) Clustering of the mouse retina bipolar cells by different methods. The cells are colored by replication IDs. Cells from six replicates were processed in two different batches (Bipolar1-Bipolar4 are replicates from batch1, and Bipolar5-6 are replicates from batch 2). (b) KL-divergence for measuring of batch mixing of different methods. (c) Batch effect mixing is improved over iterations in DESC. (d) KL-divergence decreases over iterations in DESC, indicating that batch effect removal is improved over iterations.

**Figure 3.**
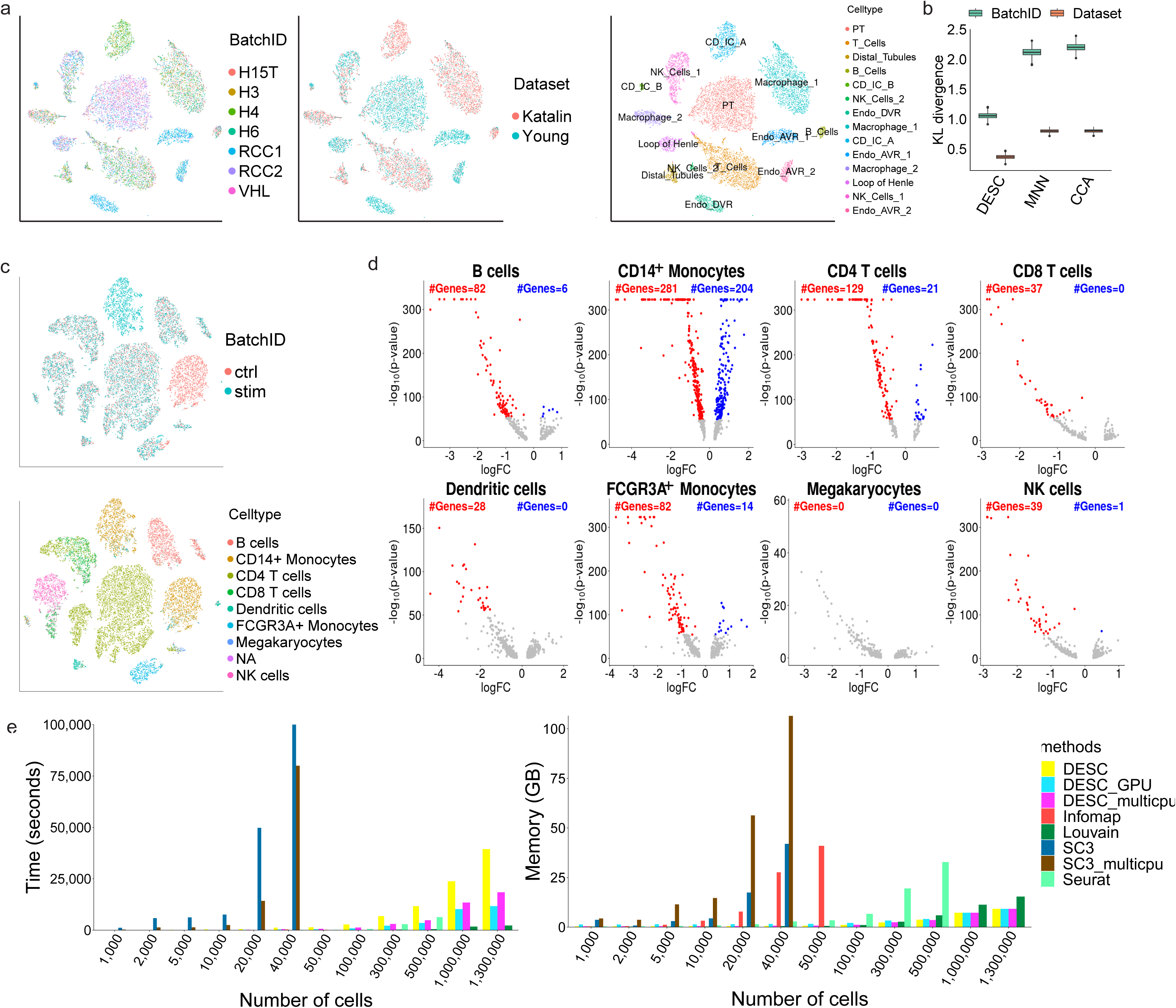
(a) DESC clustering of the human kidney data. Cell types were determined based on known marker genes. Endo_AVR: Endothelial Ascending Vasa Recta; Endo_DVR: Endothelial Descending Vasa Recta; CD-IC: Collecting Duct Intercalated Cell; NK: Natural Killer; PT: Proximal Tubule. (b) KL-divergence for measuring batch mixing of different methods for the human kidney data. (c) DESC clustering of the PBMC data. Cell types were based on assignment reported in the original paper. (d) Volcano plot of differential expression analysis between control and stimulus conditions for each cell type. Highlighted are genes with FDR adjusted p-value<10^-50^. CD14^+^ monocytes has the most number of differentially expressed genes compared to the other cell types. (e) Comparison of memory usage and running time of each method for datasets with various numbers of cells, where the cells were randomly sampled from the 1.3 million mouse brain dataset.

We also assessed the performance of DESC on data with complex batch effect generated from multiple subjects using the same platform but in different labs. Such complex batch effect is common in human studies because logistical constraints mandate that data from different subjects be generated at different times and perhaps in different labs, which result in complex batch effects that are challenging to address. To examine the robustness of DESC in the presence of this complex batch effect, we analyzed scRNA-seq data obtained from seven human kidneys (**Supplementary Note 4**). This dataset includes 8,544 cells, derived from four healthy kidneys, generated by us using 10X, and 7,149 cells obtained from the normal part of kidneys in three patients with kidney tumor^13^, also generated by 10X, but in a different lab. **Figs. 3a and b** show that DESC removed batch effect, with the seven biological samples and the two different datasets randomly mixed. The KL-divergence is lower for DESC than for CCA and MNN (**Fig. 3d**), indicating that DESC is more effective in removing batch effect both at the subject level and dataset level.

The kidneys and the immune system are closely linked. It has been shown that the accumulation of natural killer (NK) cells promotes chronic kidney inflammation and contributes to kidney fibrosis^14^. T cells, which have a well-described role in renal injury, are involved in renal fibrosis^15^. Previous studies have shown that NK cells play a role in the regulation of the adaptive immune response and stimulate or inhibit T cell responses^16^. Better understanding of how different components of the immune system mediate kidney disease requires a clear separation of NK and T cells. **Fig. 3c and Supplementary Fig. 8** show that both DESC and MNN identified T cells and NK cells as separate clusters; however, CCA mixed some of the NK cells with T cells, possibly due to overcorrection of true biological variations. These results indicate that DESC not only removed technical batch effect more effectively than CCA and MNN, but also maintained true biological variations among closely related immune cells.

To further demonstrate that DESC preserves true biological variations, we considered an even more complex situation in which technical batches were completely confounded with biological variations. This is inevitable in disease studies where tissues are processed immediately to maintain cell viability resulting in the preparation of normal and diseased samples in different batches. For data generated in such complex settings, it is desirable to remove technical batch effect while maintaining true biological variations between normal and diseased samples so that disease specific subpopulations can be identified. We analyzed a dataset generated by 10X that includes 24,679 human PBMCs from eight patients with lupus^17^ (**Supplementary Note 5**). The cells were split into a control group and a matched group stimulated with INF-β, which leads to a drastic but highly cell type-specific response. This dataset is extremely challenging because removal of technical batch effect is complicated by the presence of biological differences, both between cell types under the same condition and between different conditions.

**Fig. 3c** shows that DESC randomly mixed cells between the control and the stimulus conditions for all cell types except CD14^+^ monocytes. Differential expression (DE) analysis revealed a drastic change in gene expression after INF-β stimulation for CD14^+^ monocytes (**Fig. 3d**); the number of DE genes and the magnitude of DE, measured by p-value and fold-change, are several orders more pronounced than the other cell types. This is consistent with previous studies showing CD14^+^ monocytes with a more drastic gene expression change than B cells, dendritic cells, and T cells after INF-β stimulation^18^, ^19^. These results suggest that DESC is able to remove technical batch effect and maintain true biological variations induced by INF-β. MNN also preserved the biological difference between the control and the INF-β stimulated CD14^+^ monocytes, but the NK cells are less well separated from CD8 T cells (**Supplementary Fig. 15a**). CCA masked the biological difference between the control and the INF-β stimulated CD14^+^ monocytes indicating that it might have overcorrected batch effect (**Supplementary Fig. 15a**).

In summary, we have developed a deep learning algorithm that clusters scRNA-seq data by iteratively optimizing a clustering objective function with a self-training target distribution. DESC’s memory usage and running time increase linearly with the number of cells, thus making it scalable to large datasets (**Fig. 3e**). DESC can further speed up computation by GPUs. We analyzed a mouse brain dataset with 1.3 million cells generated by 10X, which only took about 3.5 hours with one NVIDIA TITAN Xp GPU (**Supplementary Note 6**). Compared to existing scRNA-seq clustering methods DESC improves clustering by iteratively learning cluster-specific gene expression features from cells clustered with high confidence. This iterative clustering also removes batch effect and maintains true biological differences between clusters. As the growth of single-cell studies increases, DESC will be a more precise tool for clustering of large datasets.

## ACKNOWLEDGEMENTS

This work was supported by the following funding: NIH R01GM108600, R01GM125301, and R01HL113147 (to M.L.); NIH R01DK076077 (to K.S.).

## AUTHOR CONTRIBUTIONS

This study was conceived of and led by M.L. and H.G.. X.L., H.G., and M.L. designed the model and algorithm. X.L. implemented the DESC software and led the data analysis with input from M.L., H.G., and J.Z.. Y.L. helped with software development and testing. K.S. and J.P. generated the human kidney scRNA-seq data, and provided input on the kidney data analysis. D.S. provided input on the mouse retina scRNA-seq data analysis. M.L., X.L., and H.G. wrote the paper with feedback from Y.L., J.P., J.Z., D.S., and K.S..

## COMPETING FINANCIAL INTERESETS STATEMENT

The authors declare no competing interests.

## ONLINE METHODS

### The DESC algorithm

Analysis of scRNA-seq data often involves clustering of cells into different clusters and selection of highly variable genes for cell clustering. As these are closely related, it is desirable to use a data driven approach to cluster cells and select genes simultaneously. This problem shares similarity with pattern recognition, in which clear gains have resulted from joint consideration of the classification and feature selection problems by deep learning. However, for scRNA-seq data, a challenge is that we cannot train deep neural network with labeled data as cell type labels are typically unknown. To solve this problem, we take inspiration from recent work on unsupervised deep embedding for clustering analysis^20^, in which we iteratively refine clusters with an auxiliary target distribution derived from the current soft cluster assignment. This process gradually improves clustering as well as feature representation.

#### Overview of DESC

The DESC procedure starts with parameter initialization, in which a stacked autoencoder is used for pretraining and learning low-dimensional representation of the input gene expression matrix. The corresponding encoder is then added to the iterative clustering neural network. The cluster centers are initialized by the Louvain’s clustering algorithm^5^, which aims to optimize modularity for community detection. This clustering returns data in a feature space that allows us to obtain centroids in the initial stage of the iterative clustering. Below, we describe each component of the DESC procedure in detail.

#### Parameter initialization by stacked autoencoder

Let *X ∈ R*^*n×p*^ be the gene expression matrix obtained from a scRNA-seq experiment, in which rows correspond to cells and columns correspond to genes. Due to sparsity and high-dimensionality of scRNA-seq data, to perform clustering, it is necessary to transform the data from high dimensional space *R*^*p*^ to a lower dimensional space *R*^*d*^ in which *d ≪ p*. Traditional dimension-reduction techniques such as principal component analysis, operate on a shallow linear embedded space, and thus have limited ability to represent the data. To better represent the data, we perform feature transformation by a stacked autoencoder, which have been shown to produce well-separated representations on real datasets.

The stacked autoencoder network is initialized layer by layer with each layer being an autoencoder trained to reconstruct the previous layer’s output. After greedy layer-wise training, all encoder layers are concatenated, followed by all decoder layers, in reverse layer-wise training order. The resulting autoencoder is then fine-tuned to minimize reconstruction loss. The final result is a multilayer autoencoder with a bottleneck layer in the middle. After fine tuning, the decoder layers are discarded, and the encoder layers are used as the initial mapping between the original data space and the dimension-reduced feature space, as shown in **Fig. 1a**.

Since the number of true clusters for a scRNA-seq dataset is typically unknown, we apply the Louvain’s method, a graph-based method that has been shown to excel over other clustering methods, on the feature space *Z* obtained from the bottleneck layer. This analysis returns the number of clusters, denoted by *K*, and the corresponding cluster centroids {*μ*_*j*_: *j* = 1, …, *K*}, which will be used as the initial clustering for DESC.

#### Iterative clustering

After cluster initialization, we improve the clustering using an unsupervised algorithm that alternates between two steps until convergence. In the first step, we compute a soft assignment of each cell between the embedded points and the cluster centroids. Following van der Maaten & Hinton^21^, we use the Student’s *t*-distribution as a kernel to measure the similarity between embedded point *z*_*i*_ for cell *i* and centroid *μ*_*j*_ for cluster *j*,

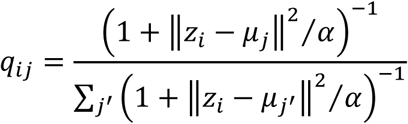

where *z*_*i*_ = *f*_*W*_(*x*_*i*_) ∈ *Z* corresponds to *x*_*i*_ ∈ *X* after embedding, α is the degree of freedom of the Student’s *t*-distribution.

In the second step, we refine the clusters by learning from cells with high confidence cluster assignments with the help of an auxiliary target distribution. Specifically, we define the objective function as a Kullback-Leibler (KL) divergence loss between the soft cell assignments *q*_*i*_ and the auxiliary distribution *p*_*i*_ for cell *i* as

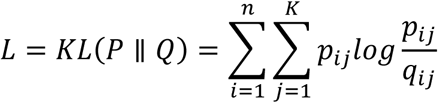

where the auxiliary distribution *P* is defined as

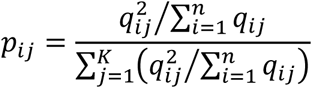

The encoder is fine-tuned by minimizing *L*. The above definition of the auxiliary distribution *P* can improve cluster purity by putting more emphasis on cells assigned with high confidence. Given that the target distribution *P* is defined by *Q*, minimizing *L* implies a form of self-training. Also, *p*_*ij*_ gives the probability of cell *i* that belongs to cluster *j*, and this probability can be used to measure the confidence of cluster assignment for each cell. Because α is insensitive to the clustering result, we let α = 1 for all datasets by default.

#### Optimization of the KL divergence loss

We jointly optimize the cluster centers {*μ*_*j*_: *j* = 1, …, *K*} and the deep neural network parameters using stochastic gradient descent. The gradients of *L* with respect to feature space embedding of each data point *z*_*i*_ and each cluster center *μ*_*j*_ are

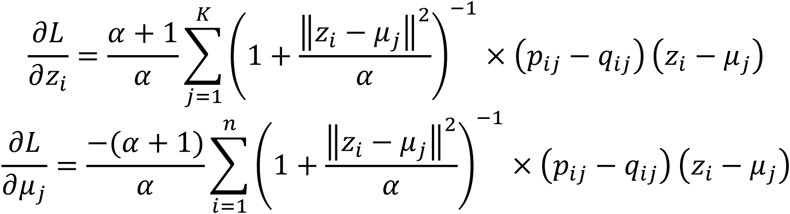

These gradients are then passed down to the deep neural network and used in standard backpropagation to compute the deep neural network’s parameter gradient. We use Keras to train our model. During each iteration i.e. when loss is not decreasing or the epoch number threshold is reached, we update the auxiliary distribution *P*, and optimize cluster centers and encoder parameters with the new *P*. This iterative procedure is stopped when less than *tol*% of cells change cluster assignment between two consecutive iterations. We let *tol* = 0.5 by default.

#### Architecture of the deep neural network in DESC

Depending on the number of cells in the dataset, we suggest different numbers of hidden layers and different numbers of nodes in the encoder. **Supplementary Table 2** gives the default numbers of hidden layers and nodes in DESC.

DESC allows users to specify their own numbers of hidden layers and nodes. We recommend using more hidden layers and more nodes per layer for datasets with more cells so that the complexity of the data can be captured by the deep neural network. We use ReLU as the activation function except for the last hidden layer and last decoder layer, in which tanh is used as the activation function. The reason why we use tanh is that we must guarantee the values in feature representation and output of decoder range from negative to positive. The default hyperparameters for the autoencoder are listed in **Supplementary Table 3**.

### Data normalization and gene selection

The normalization involves two steps. In the first step, cell level normalization is performed, in which the UMI count for each gene in each cell is divided by the total number of UMIs in the cell, and then transformed to a natural log scale. In the second step, gene level normalization is performed in which the cell level normalized values for each gene are standardized by subtracting the mean across all cells and divided by the standard deviation across all cells for the given gene. Highly variable genes are selected using the filter_genes_dispersion function from the Scanpy package^22^ (https://github.com/theislab/scanpy).

### Evaluation metric for clustering

For published datasets in which the reference cell type labels are known, we use ARI to compare the performance of different clustering algorithms. Larger values of ARI indicate higher accuracy in clustering. The ARI can be used to calculate similarity between the clustering labels obtained from a clustering algorithm and the reference cluster labels. Given a set of *n* cells and two sets of clustering labels of these cells, the overlap between the two sets of clustering labels can be summarized in a contingency table, in which each entry denotes the number of cells in common between the two sets of clustering labels. Specifically, the ARI is calculated as

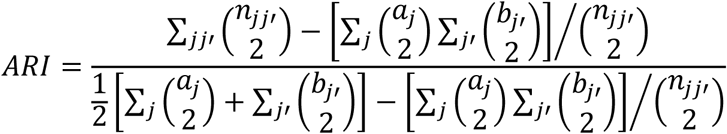

where *n*_*jj’*_ is the number of cells assigned to cluster *j* based on the reference cluster labels, and cluster *j’* based on clustering labels obtained from a clustering algorithm, *a*_*j*_is the number of cells assigned to cluster *j* in the reference set, and *b*_*j’*_ is the number of cells assigned to cluster *j’* by the clustering algorithm.

### Evaluation metric for batch effect removal

We use KL-divergence to evaluate the performance of various single-cell clustering algorithms for batch effect removal i.e., how randomly are cells from different batches mixed together within each cluster. The KL-divergence of batch mixing for *B* different batches is calculated as

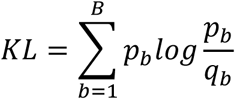

where *q*_*b*_ is the proportion of cells from batch *b* among all cells, and *p*_*b*_ is the proportion of cells from batch *b* in a given region based on results from a clustering algorithm, with 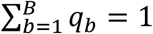 and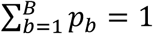. We calculate the KL divergence of batch mixing on the first two components of the t-SNE coordinates, by using regional mixing KL divergence defined above at the location of 100 randomly chosen cells from all batches. The regional proportion of cells from each batch is calculated based on the set of 120 nearest neighboring cells from each randomly chosen cell. The final KL divergence is then calculated as the average of the regional KL divergence. We repeated this procedure for 500 iterations with different randomly chosen cells to generate box plots of the final KL divergence. Smaller final KL divergence indicates better batch mixing i.e., more effective in batch effect removal.

### Datasets

We analyzed multiple scRNA-seq datasets. Publicly available data were acquired from the access numbers provided by the original publications. The human kidney dataset generated by us is available in Supplementary Data.

#### Benchmarking dataset

The Genotype-Tissue Expression (GTEx) v7 dataset^10^ was downloaded from the GTEx data portal (https://gtexportal.org/home/datasets). This dataset includes 11,688 human RNA-seq samples from 30 tissues. Because the tissue origin is known, we treat this dataset as the benchmarking dataset in which the tissue origin is used as the true cluster label.

#### Drosohpila dataset

The data were generated by Croset et al.^11^ in which 10,286 cells were generated using Drop-seq from the midbrain of drosophila.

#### Mouse retina dataset

The data were generated by Shekhar et al.^12^ in which 23,494 bipolar cells were generated using Drop-seq from retinas of six mice processed in two experimental batches. This dataset allows us to examine batch effect at the subject level.

#### Human kidney datasets

The first set of data was generated by us using 10X. This dataset includes 8,544 cells from kidneys in four healthy human subjects. The second set of data was generated by Young et al.^13^, also using 10X. This dataset includes 7,149 cells from the normal part of the kidneys in three human subjects that have kidney tumors. These two datasets were combined in our analysis, which allow us to examine batch effect, both at the subject level and at the dataset level.

#### Human PBMC dataset

The data were generated by Kang et al.^17^ in which 24,679 PBMC cells were obtained and processed from eight patients with lupus using 10X. These cells were split into two groups: one stimulated with interferon-beta (INF-β) and a culture-matched control. This dataset allows us to examine whether technical batch effect can be removed in the presence of true biological variations.

*1.3 million brain cells from E18 mice.* This dataset was downloaded from the 10X Genomics website (https://support.10xgenomics.com/single-cell-gene-expression/datasets/1.3.0/1Mneurons). It includes 1,306,127 cells from cortex, hippocampus and subventricular zone of two E18 C57BL/6 mice.

A complete list of the datasets analyzed in this paper is provided in **Supplementary Table 1**.

### Software availability

An open-source implementation of the DESC algorithm can be downloaded from https://eleozzr.github.io/desc/.

