## Supplementary material for "Deep learning enables accurate clustering and batch effect removal in single-cell RNA-seq analysis"

**Supplementary Table 1. Datasets analyzed in this paper**

**Supplementary Table 2. The numbers of hidden layers and nodes in the encoder**

**Supplementary Table 3. Default hyperparameters of the autoencoder**

**Supplementary Note 1. Analysis of the GTEx data**

**Supplementary Note 2. Analysis of the Drosophila midbrain data**

**Supplementary Note 3. Analysis of the mouse retina data**

**Supplementary Note 4. Analysis of the human kidney data**

**Supplementary Note 5. Analysis of the human PBMC data**

**Supplementary Note 6. Analysis of the mouse brain data with 1.3 million cells**

**Supplementary Note 7. Computing time and memory usage**

**Supplementary Table 1. Datasets analyzed in this paper**

| Species | Tissue | Data source | No. of subjects | Sample size | Protocol |
|---|---|---|---|---|---|
| Human | 30 tissues | GTEx | - | 11,687 samples | Bulk RNA-seq |
| Drosophila | Mid brain | Croset et al. (2018) | 8 | 10,286 cells | Drop-seq |
| Mouse | Retina | Shekhar et al. (2016) | 6 | 27,499 cells | Drop-seq |
| Human | Kidney | Generated by us | 4 | 8,544 cells | 10X |
| Human | Kidney | Young et al. (2018) | 3 | 7,149 cells | 10X |
| Human | PBMC | Kang et al. (2018) | 2 | 24,679 cells | 10X |
| Mouse | Brain | 10X website | 2 | 1,306,127 cells | 10X |

**Supplementary Table 2. The numbers of hidden layers and nodes in the encoder**

| No. of Cells | No. of hidden layers | No. of nodes in the 1st hidden layer | No. of nodes in the 2nd hidden layer |
|---|---|---|---|
| >20,000 | 2 | 128 (or larger) | 32 |
| (10,000,20,000] | 2 | 64 | 32 |
| (5,000,10,000] | 2 | 32 | 16 |
| (2,000,5,000] | 1 | 128 | 0 |
| (500,2,000] | 1 | 64 | 0 |
| <500 | 1 | 16 | 0 |

**Supplementary Table 3. Default hyperparameters of the autoencoder**

| Parameter | Default value |
|---|---|
| Activation function | ReLU or Tanh |
| Kernel initializer | glorot_uniform |
| Dropout rate | 0.2 |
| Optimizer | Stochastic gradient descent |
| Learning rate | 0.01 |
| Batch Size | 256 |
| No. of epochs | 300 |

### Supplementary Note 1: analysis of the GTEx data

The Genotype-Tissue Expression (GTEx) data were downloaded from the GTEx Portal website (https://gtexportal.org/home/datasets). We downloaded data in the current release (v7) (https://storage.googleapis.com/gtex_analysis_v7/rna_seq_data/GTEx_Analysis_2016-01-15_v7_RNASeQCv1.1.8_gene_reads.gct.gz), which include 11,688 RNA-seq samples and 56,202 genes from 30 unique human tissues.

Gene filtering criteria: a gene was eliminated if the number of samples expressing this gene is <20.

Data processing: 1) gene expression levels for each sample was normalized using the "scanpy.api.normalize_per_cell" function in scanpy with default parameters; 2) top 1,000 highly variable genes were selected using the "scanpy.api.pp.filter_genes_dispersion" function in scanpy, which is the same as the function 'FindVariableGenes' in Seurat; 3) normalized gene expression for the selected top 1,000 highly variable genes was transformed using log(1+x) transformation with natural logarithm; 4) the expression value was further standardized to a z-score, and the standardized gene expression values were used as input for DESC.

After the above filtering and data processing, there were 11,687 samples×1,000 highly variable genes remained in DESC analysis. The same highly variable genes were used as input for Louvain's method. For SC3, we used the default parameter values of function sc3 in R package SC3. For Infomap, we used the default parameter values of function prefilterGenes in R package SINCERA.

DESC analysis: We used two hidden layers for the encoder model with 64 nodes in the first hidden layer, and 32 nodes in the second hidden layer. Other parameters were set at default values. The final model is 1000-64-32-64-1000.

Evaluation metrics for clustering: In addition to ARI, a metric described in the main text, we also evaluated the performance of different clustering algorithms using other metrics, including normalized mutual information (NMI) and purity, which are calculated as the following.

NMI: Cluster labels in the reference set defines the probability distribution $P_R(j) = \frac{n_j}{n}$, where $n_j$ is the number of cells in cluster $j$ of the reference set, and $n$ is the total number of cells in the dataset. A clustering algorithm also determines a probability distribution $P_C(j') = \frac{n_{j'}}{n}$, where $n_{j'}$ is the number of cells in cluster $j'$ based on the clustering algorithm. For the probability distributions, $P_R$ and $P_C$, their mutual information is

$$I(P_R, P_C) = \sum_{j,j'} R(j,j') log \frac{R(j,j')}{P_R(j) P_C(j')}$$

where $R(j,j') = \frac{n_{jj'}}{n}$ is defined by the joint probability distribution. The NMI is calculated as

$$NMI = \frac{I(P_R, P_C)}{\sqrt{H(P_R) H(P_C)}}$$

where $H(P_R) = \sum_j P_R(j) log\big(P_R(j)\big)$, and $H(P_C) = \sum_{j'} P_C(j') log\big(P_C(j')\big)$.

Purity: Purity is the percent of the total number of cells that are classified correctly. It is calculated as

$$purity = \frac{1}{n}\sum_{j=1}^{K} \max_{j'} |C_j \cap T_{j'}|$$

where $n$ is the total number of cells in the dataset, $K$ is the total number of cell types based on the reference cluster labels, $C_j$ is a cluster in the reference data, and $T_{j'}$ is the cluster which shares the most number of cells with cluster $C_j$.

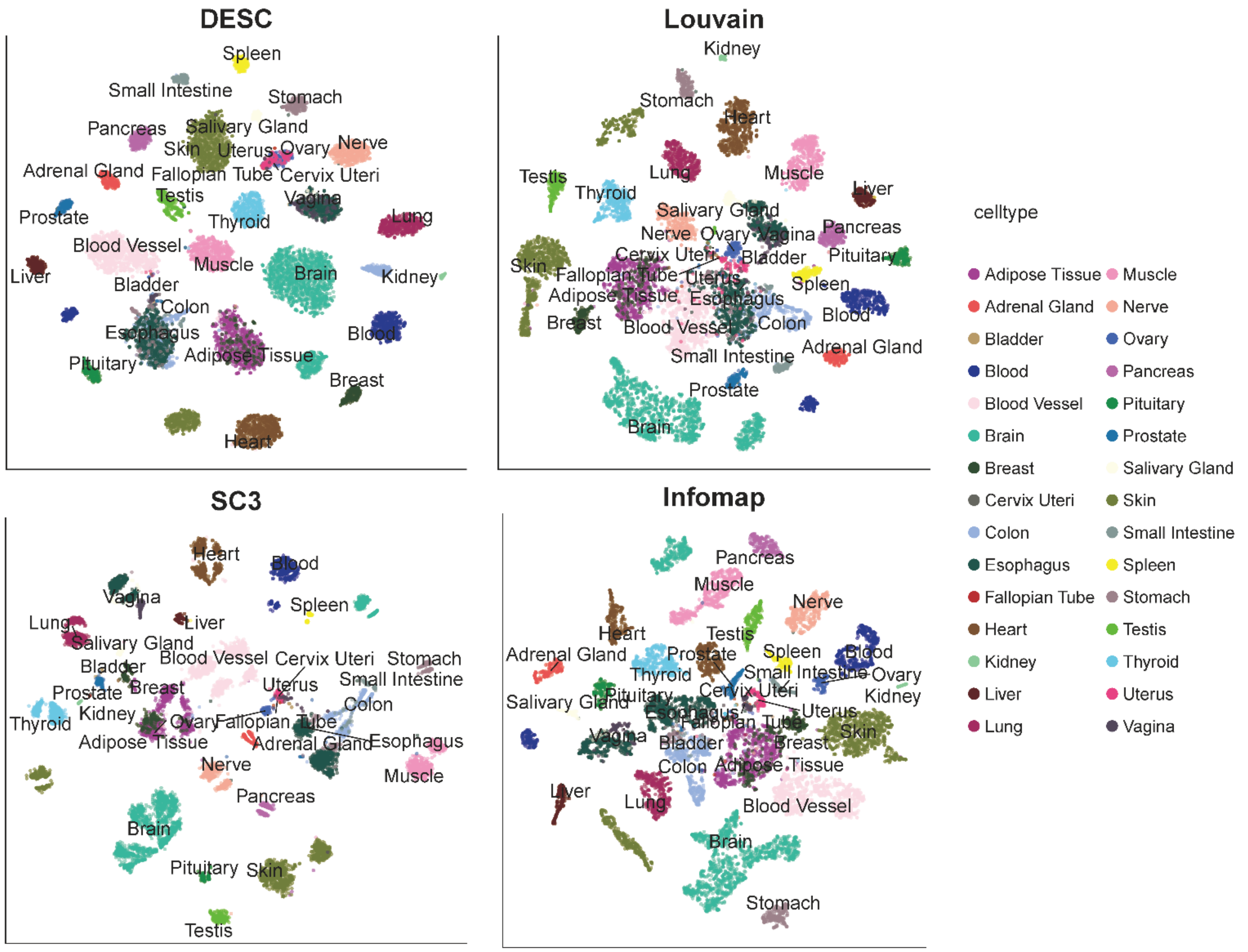


**Supplementary Fig 1.** T-SNE plots for clustering results of DESC, Louvain's method, SC3 and Infomap on the GTEx data. Displayed are the best clustering results for each method based on ARIs. The resolution was set at 0.4 for DESC, and 0.2 for Louvain's method implemented in scanpy. For SC3, the number of clusters was set at 28. For Infomap, the number of PCs was set at 50.

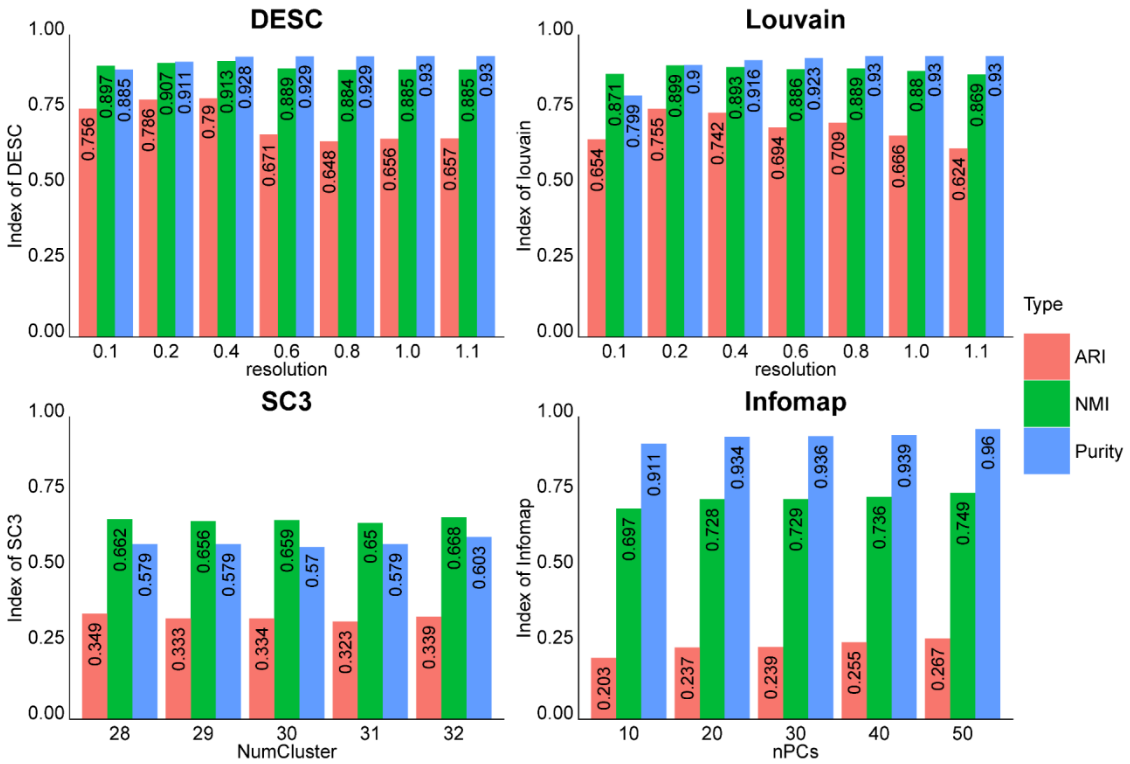


**Supplementary Fig 2.** Clustering evaluation metrics for DESC, Louvain's method, SC3 and Infomap on the GTEx data. The original tissue origin of each sample was treated as the true cluster label when calculating the clustering evaluation metrics. For each clustering algorithm, we evaluated its performance with different resolutions (for DESC and Louvain's method), different numbers of clusters (for SC3), and different numbers of PCs (for Infomap).

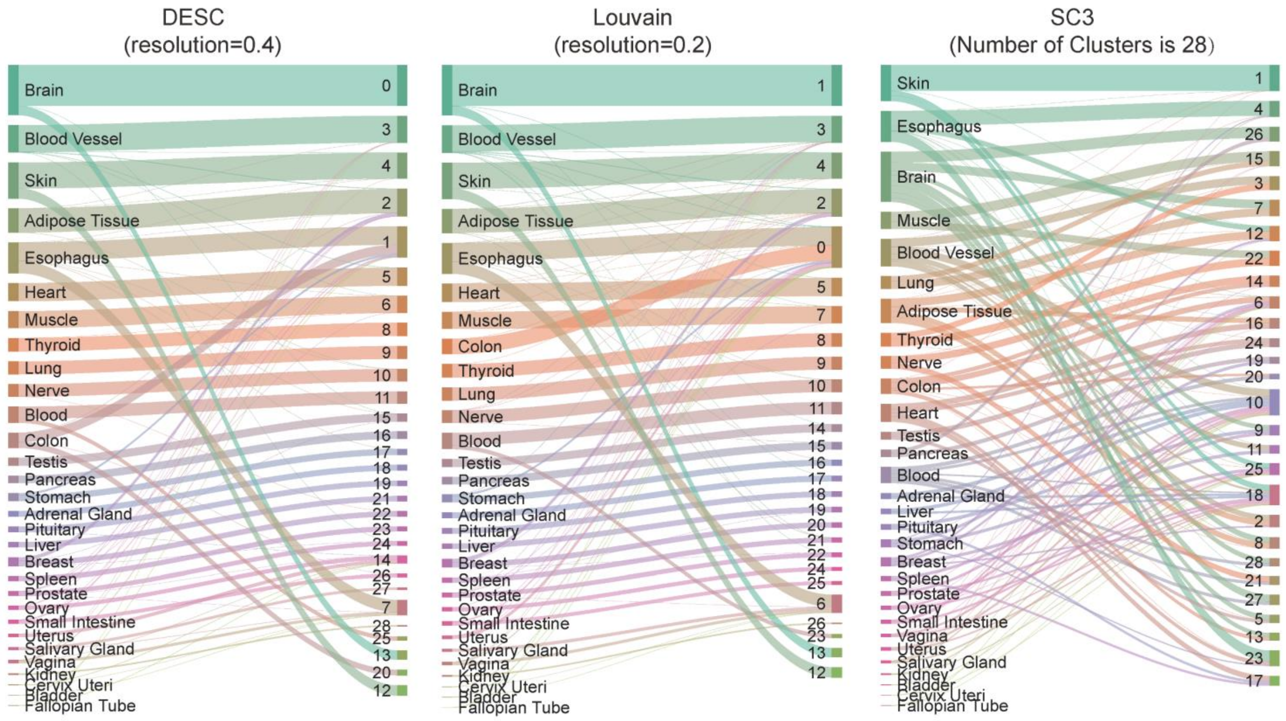


**Supplementary Fig 3.** Sankey diagrams of the DESC, Louvain's and SC3 clustering results on the GTEx data. The resolution was set at 0.4 for DESC and 0.2 Louvain's clustering. The number of clusters for SC3 was set at 28. We did not include Infomap in this visualization because Infomap tends to produce many small clusters, typicall more than 100, across all parameter settings we evaluated.

**Supplementary Note 2: analysis of the Drosophila mid brain data**

This dataset was generated by Croset et al. (2018) Cellular diversity in the Drosophila midbrain revealed by single-cell transcriptomics. Elife 7. pii:e34550.

The raw gene expression count matrix, which includes 10,286 cells and 10,934 genes was download from https://elifesciences.org/download/aHR0cHM6Ly9jZG4uZWxpZmVzY2llbmNlcy5vcmcvYXJ0aWNsZXMvMzQ1NTAvZWxpZmUtMzQ1NTAtZmlnMS1kYXRhMS12Mi56aXA=/elife-34550-fig1-data1-v2.zip?_hash=bJMTSD0Bed%2FkJDq3MxXZbUEoxokD1Fnfa36O2P2WnQs%3D.

The t-SNE coordinates and metadata for Figure 1 reported in the paper were downloaded from http://scope.aertslab.org/#/d5c805ab-fb1e-4ccf-ae81-a53bbab1a4d2/Waddell_CentralBrain_10k.loom/gene.

Cell filtering criteria: 1) We didn't filter out any cells because the downloaded data were already prefiltered.

Gene filtering criteria: a gene was eliminated if the number of cells expressing this gene is <10.

Data processing: 1) gene expression levels for each cell was normalized using the "scanpy.api.normalize_per_cell" function in scanpy with counts_per_cell_after =10,000 ; 2) top 1,000 highly variable genes were selected using the "scanpy.api.pp.filter_genes_dispersion" function in scanpy; 3) normalized gene expression for the selected top 1,000 highly variable genes was then transformed using log(1+x) transformation with natural logarithm; 4) the expression value is further standardized to a z-score, and the standardized gene expression values were used as input for DESC.

After the above filtering and data processing, there were 10,286 cells×1,000 highly variable genes remained in DESC analysis.

DESC analysis: We used two hidden layers for encoder with 64 nodes in the first hidden layer, and 32 nodes in the second hidden layer. Other parameters were set as default values. The final model is 1000-64-32-64-1000.

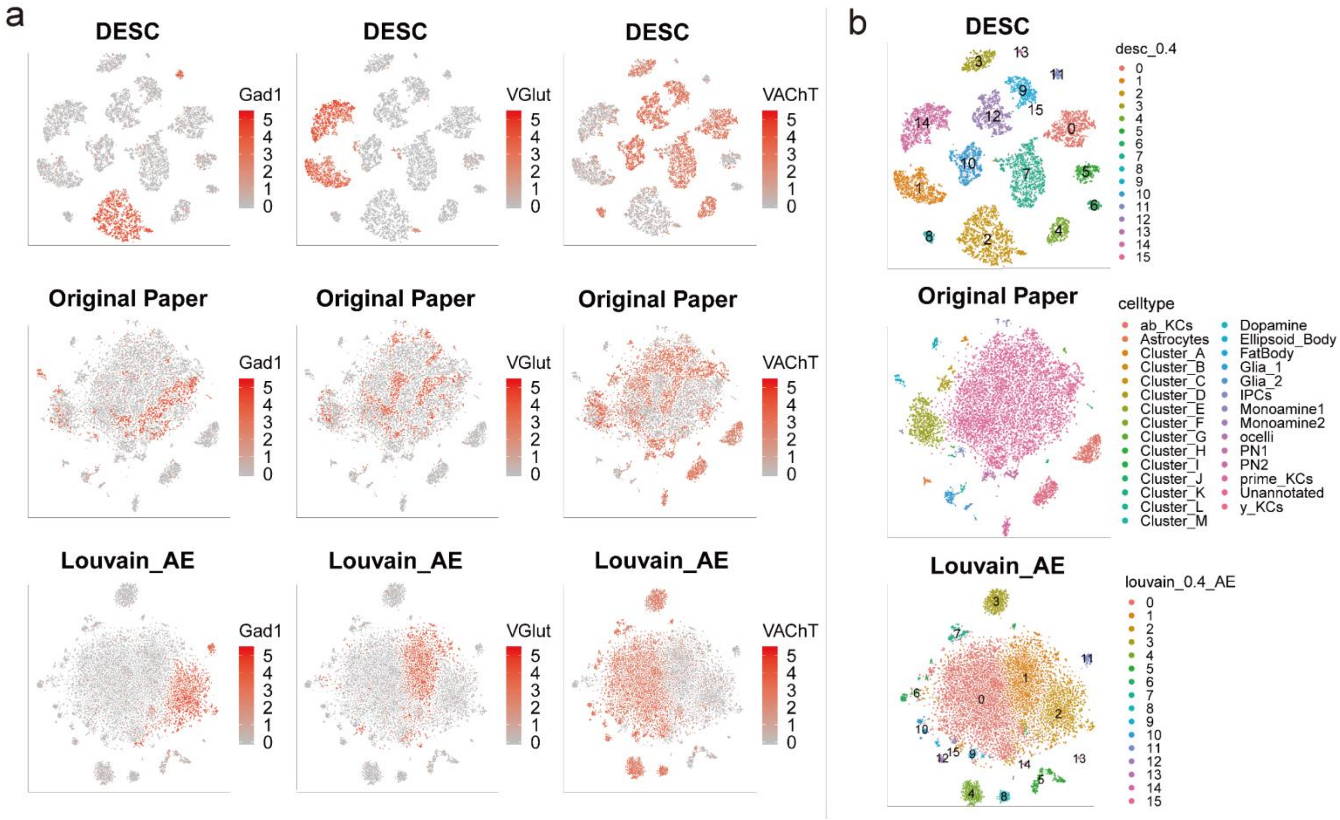


**Supplementary Fig 4. (a)** Gene expression feature plot of the Drosophila dataset for Gad1 (marker gene for cholinergic neurons), VGlut (marker gene for glutamatergic neurons), and VAChT (marker gene for GABAergic neurons). Top panel is the result of DESC when resolution=0.4, middle panel is the result of the original paper, and bottom panel is the result of Louvain's methods using the low dimensional representation learned from the autoencoder in DESC as input. **(b)** The t-SNE plots of DESC, original paper, and Louvain with low dimensional representation learned from the autoencoder as input, where the cells were colored by the respective cluster ID or cell type.

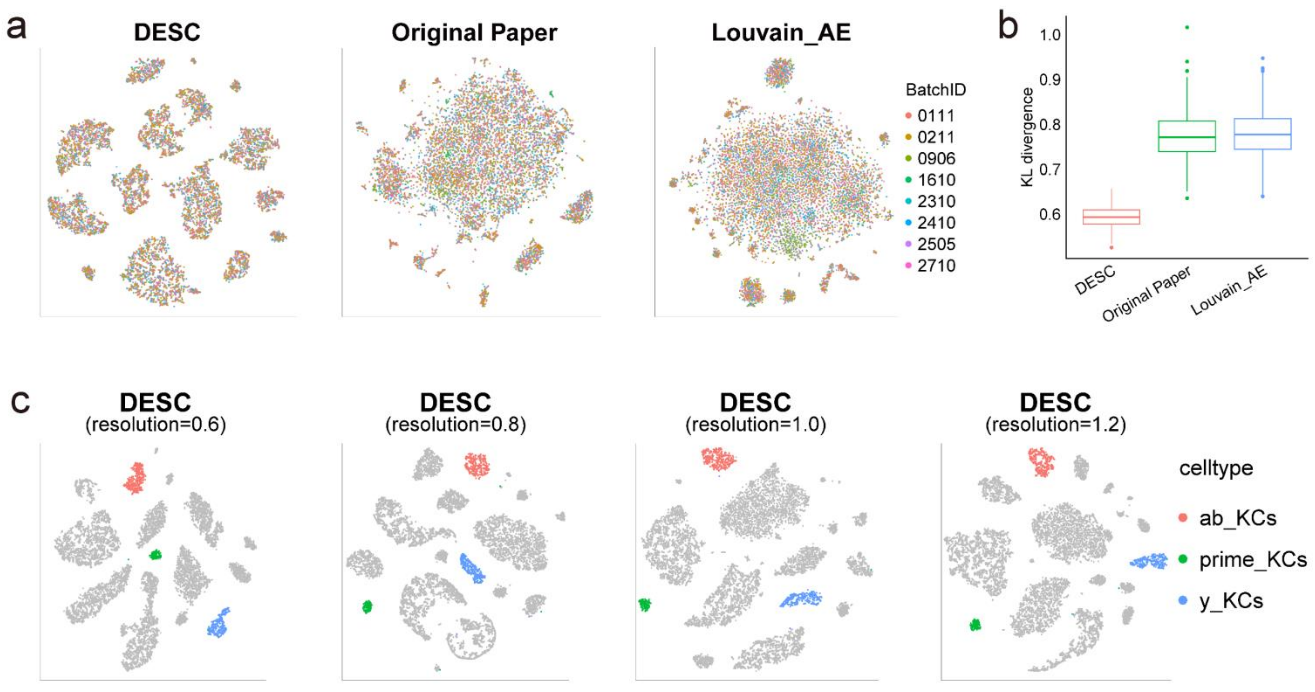


**Supplementary Fig 5. (a)** Drosophila dataset t-SNE plots of the DESC clustering (resolution = 0.4) (left), Louvain's clustering reported in the original paper (middle), and Louvain's clustering based on the representation of autoencoder (right), where the cells were colored by batch IDs. **(b)** The KL-divergence of that measures the degree of batch effect removal for the three clustering results. Although there is no obvious batch effect from (a), the KL-divergence index of DESC is less than the Original Paper's and Louvain's method using low-dimensional representation learned from the autoencoder. **(c)** The t-SNE plots of the DESC clustering with different resolutions (0.6, 0.8, 1.0, 1.2) in which the cells were colored by the three Kenyon cells. This result indicates that DESC is robust to the choice of resolution.

**Supplementary Note 3: analysis of the bipolar cells of mouse retina data**

This dataset was generated by Shekhar et al. (2016) Comprehensive classification of retinal bipolar neurons by single-cell transcriptomics. Cell 166(5):1308-1323.

The dataset was downloaded from https://scrnaseq-public-datasets.s3.amazonaws.com/scater-objects/shekhar.rds, which includes 27,499 cells and 13,166 genes.

Cell filtering criteria: 1) Only kept the14 main bipolar cells (RBC, BC1A, BC1B, BC2, BC3A, BC3B, BC4, BC5A, BC5B, BC5C, BC5D, BC6, BC7, BC8/9), which left with 23,494 cells for analysis.

Gene filtering criteria: a gene was eliminated if the number of cells expressing this gene is <10.

Data processing: 1) gene expression levels for each cell was normalized using the "scanpy.api.normalize_per_cell" function in scanpy with counts_per_cell_after =10,000; 2) top 1000 highly variable genes were selected using the "scanpy.api.pp.filter_genes_dispersion" function in scanpy; 3) normalized gene expression for the selected top 1,000 highly variable genes was then transformed using log(1+x) transformation with natural logarithm; 4) the expression value is further standardized to a z-score, and the standardized gene expression values were used as input for DESC.

After the above filtering and data processing, there were 23,494 cells×1,000 highly variable genes remained in DESC analysis.

DESC analysis: we used two hidden layers for encoder with 128 nodes in the first hidden layer, and 32 nodes in the second hidden layer. Other parameters were set default values. The final model is 1000-128-32-128-1000.

Other analyses: For CCA, we used the same number of cells, and performed analysis following Seurat's CCA tutorial (https://satijalab.org/seurat/immune_alignment.html). Specifically, we selected top 1,000 highly variable genes for each subject, pooled these genes together and then removed genes that are expressed only in a single subject. For CCA_top1000, we used the same number of cells, but the same top 1,000 highly variable genes used be DESC. For MNN, we used the same number of cells, and highly variable genes were selected using function `scanpy.api.pp.fiter_gene_ dispersion` in python module scanpy with default parameters. For MNN_top1000, we used the same number of cells and the same top 1,000 highly variable genes as DESC.

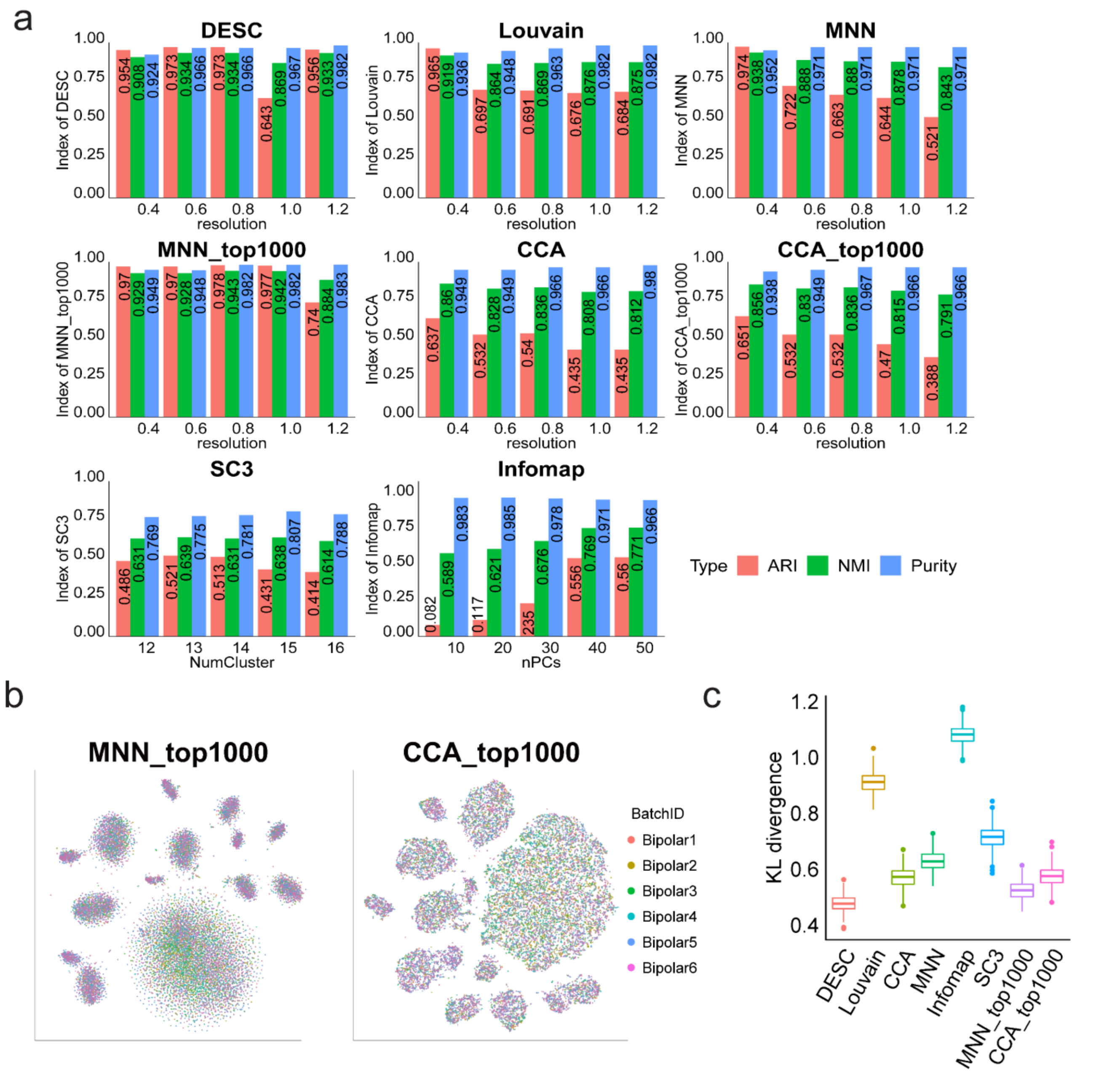


**Supplementary Fig 6. (a)** Clustering evaluation metrics for DESC, Louvain's method, SC3, Infomap, MNN, MNN_top1000, CCA, and CCA_top1000 on retinal bipolar cells. The cell type assignment reported in the original paper was treated as the true cluster label. The original paper assigned cell types using Louvain's method implemented in R package igraph with 37 significant PCs as input. For each clustering algorithm, we evaluated its performance with different resolution (for DESC, Louvain's method, MNN, MNN_top1000, CCA, CCA_top1000), different number of clusters (for SC3), and different number of PCs (for Infomap). **(b)** Cells were colored by subject IDs for MNN_top1000 and CCA_top1000. **(c)** KL-divergence for batch effect removal of different clustering algorithms.

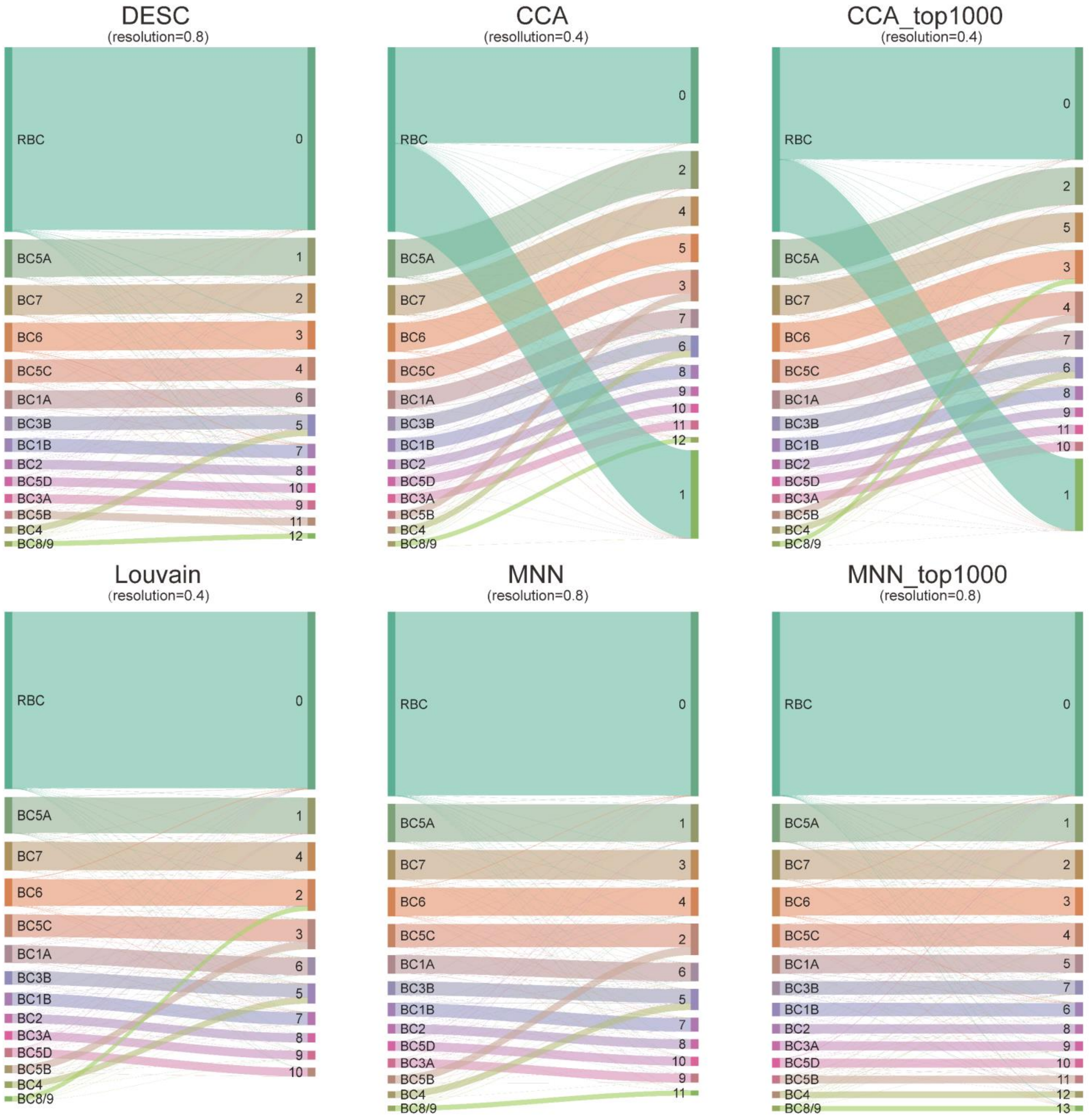


**Supplementary Fig 7.** The Sankey diagrams of different clustering algorithms on the retinal bipolar dataset. DESC, MNN, MNN_top1000 tend to cluster two similar cell type into a group, but Louvain and CCA_top1000 will gather two dissimilar cell types (BC8/9, BC6) into a group.

**Supplementary Note 4: analysis of the human kidney data**

This analysis focused on two scRNA-seq datasets generated from human kidney.

Dataset 1: The dataset includes 35,908 cells and 32,738 genes from 4 normal human kidneys, generated by Katalin Susztak's lab using 10X.

Dataset 2: The dataset was generated by Young et al. (2018) Single-cell transcriptomes from human kidneys reveal the cellular identity of renal tumors. Science 361(6402):594-599.

The dataset was download from Data S1 (http://science.sciencemag.org/highwire/filestream/713964/field_highwire_adjunct_files/4/aat1699_DataS1.gz.zip) in the Supplementary Materials of Young et al. (2018). In this analysis, we focused on normal kidney cells (total 10,621 cells), which are from VHL (2,706 cells), RCC1 (3,747 cells) and RCC2 (4,168 cells).

We combined Dataset 1 and Dataset 2 in the analysis. The resulting data include 46,529 cells, and 31,232 shared genes between these two datasets. The combined data can be downloaded from https://www.dropbox.com/s/l9zq2sge93n4ifj/human_kidney_desc_use.tar.gz?dl=0.

Cell filtering criteria: 1) eliminated cells with percentage of mitochondrial UMI counts >20%; 2) eliminated cells with gene counts <200; 3) eliminated cells with total UMI counts <1,000.

Gene filtering criteria: 1) eliminated genes if the number of cells expressing this gene is <10.

Data processing: 1) gene expression levels for each cell was normalized using the "scanpy.api.normalize_per_cell" function in scanpy with counts_per_cell_after =10,000; 2) top 1,000 highly variable genes were selected using the "scanpy.api.pp.filter_genes_dispersion" function in scanpy; 3) normalized gene expression for the selected top 1,000 highly variable genes was then transformed using log(1+x) transformation with natural logarithm; 4) the expression is further standardized to a z-score, and the standardized gene expression values were used as input for DESC.

After the above filtering and data processing, there were 15,693 cells×1,000 highly variable genes remained in DESC analysis.

DESC analysis: We used two hidden layers with 128 nodes in the first hidden layer, and 32 nodes in the second hidden layer. Other parameters are set default value. So the final model is 1000-128-32-128 -1000.

Other analyses: For CCA, we used the same number of cells as DESC, and performed analysis following Seurat's CCA tutorial (https://satijalab.org/seurat/immune_alignment.html). Specifically, we selected top 2,000 highly variable genes in each kidney, pooled these genes together and then removed the genes that are expressed in less than 5 kidneys. We required 5 kidneys to make sure that these selected genes do not express only in cells from the same lab. We also tried top 1,000 and 5,000 highly variable genes for CCA, but they were less effective in removing batch effect. For MNN, we used the same number of cells and the same top 1,000 highly variable genes as DESC.

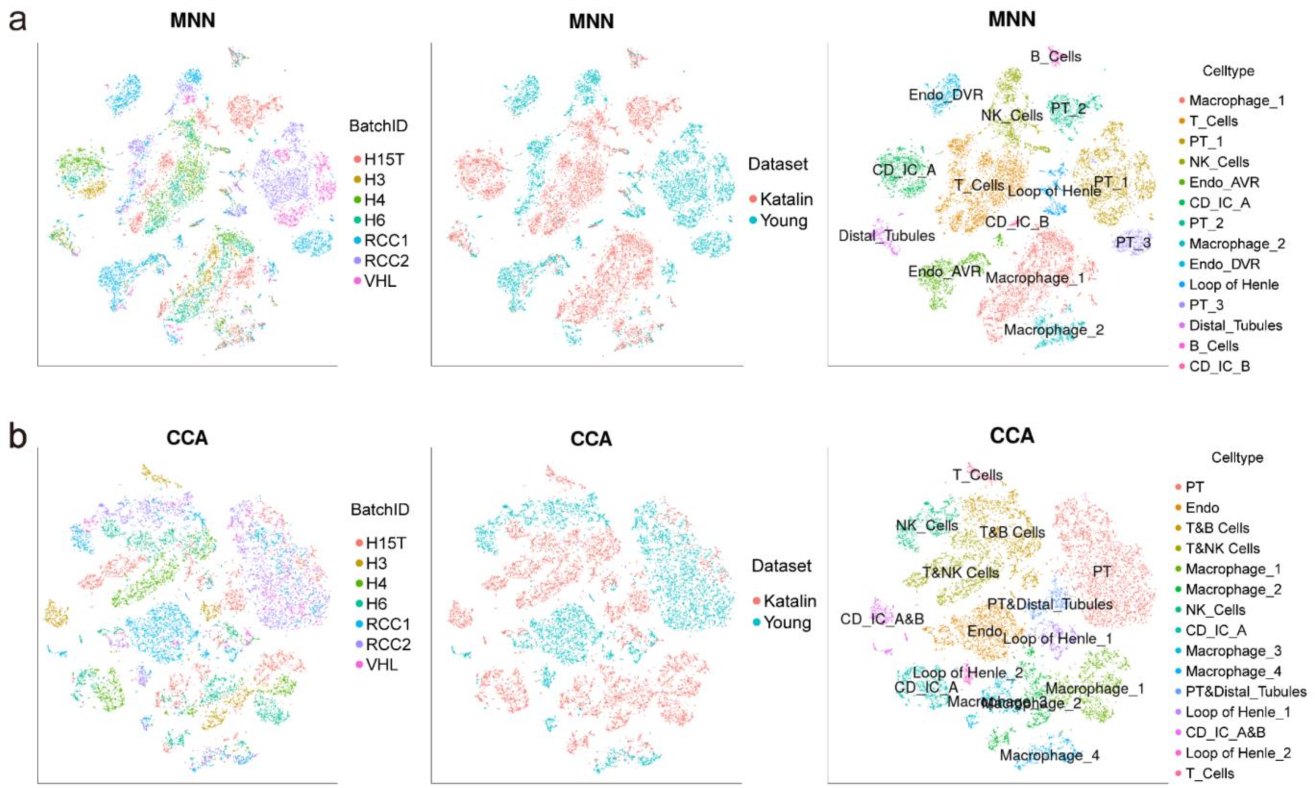


**Supplementary Fig 8. (a)** MNN and **(b)** CCA clustering results of the human kidney data. The cell types were determined based on known marker genes. Endo_AVR: Endothelial Ascending Vasa Recta; Endo_DVR: Endothelial Descending Vasa Recta; CD-IC: Collecting Duct Intercalated Cell; NK: Natural Killer; PT: Proximal Tubule.

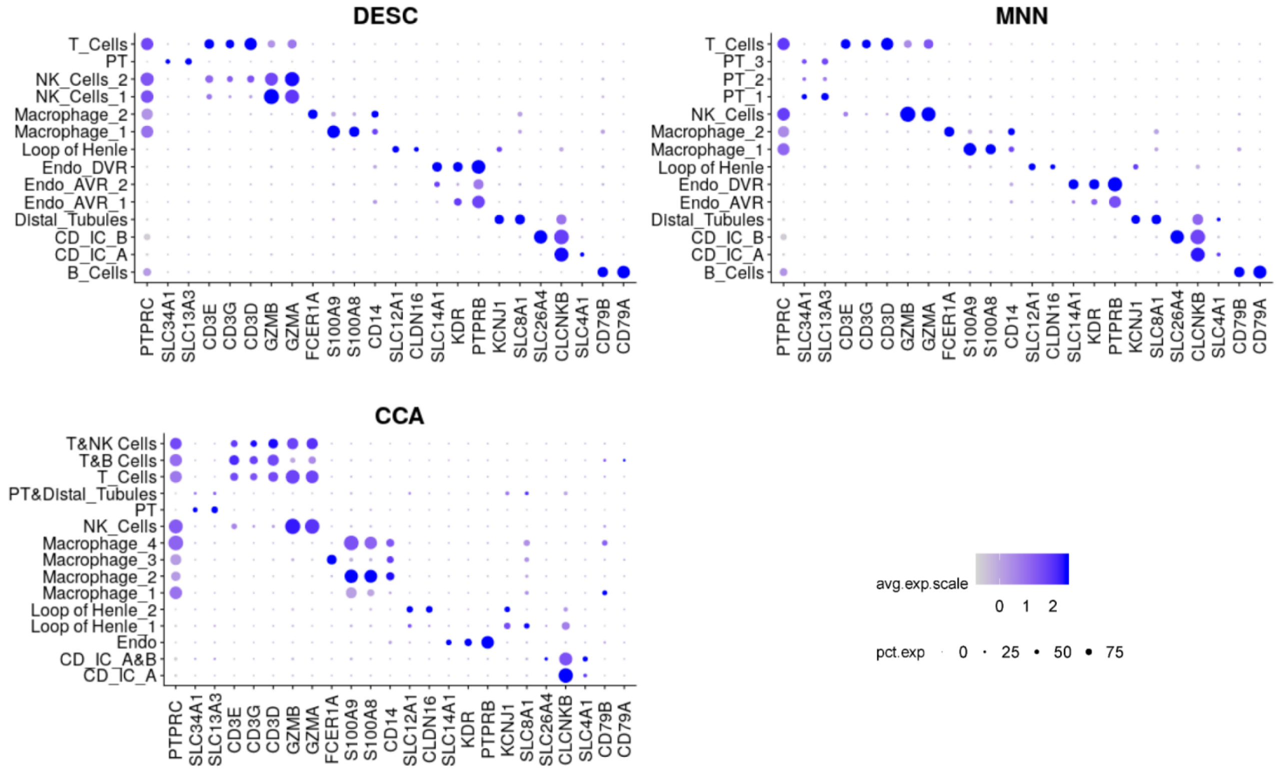


**Supplementary Fig 9.** Dot plots of known marker genes used for cell type determination for DESC, MNN, and CCA clustering results. We labeled cell types for DESC clustering using the following marker genes: SLC13A3 and SLC34A1 for PT (Proximal Tubule); CLDN16 and SLC12A for Loop of Henle; PTPRB and KDR for Endo_AVR (Endothelial Ascending Vasa Recta); PTPRB, KDR, and SLC14A1 for Endo_DVR (Endothelial Descending Vasa Recta); SLC4A1 and CLCNKB for CD_IC_A); SLC26A4 and CLCNKB for CD_IC_B; GZMA and GZMB for NK_cells; CD3D, CD3E, and CD3G for T_cells; CD14, S100A8, and S100A9 for Macrophage_1; CD14 and FCER1A for Macrophage_2; CD79A and CD79B for B_cells.

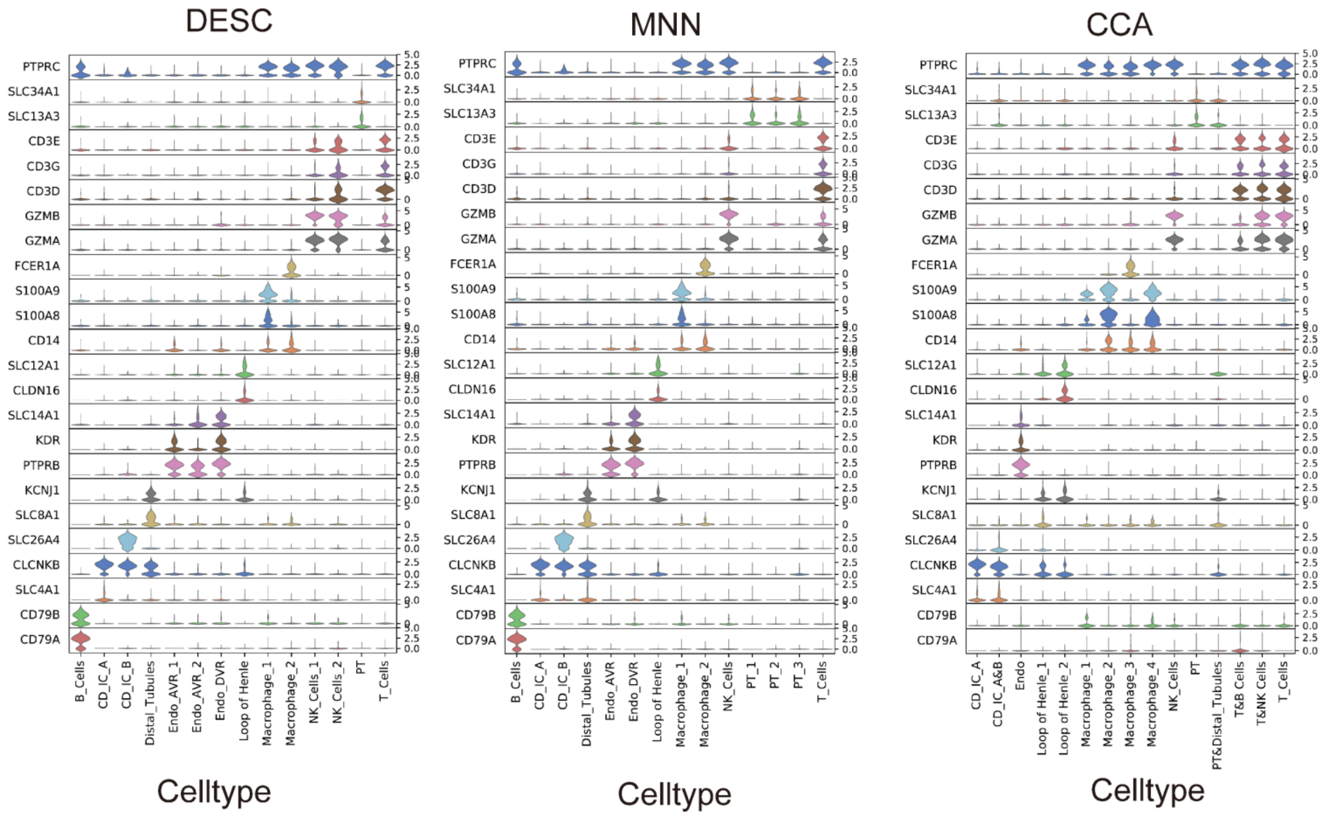


**Supplementary Fig 10.** Violin plots of known marker genes used for cell type determination for DESC, MNN, and CCA. We labeled cell types for DESC clustering using the following marker genes: SLC13A3 and SLC34A1 for PT (Proximal Tubule); CLDN16 and SLC12A for Loop of Henle; PTPRB and KDR for Endo_AVR (Endothelial Ascending Vasa Recta); PTPRB, KDR, and SLC14A1 for Endo_DVR (Endothelial Descending Vasa Recta); SLC4A1 and CLCNKB for CD_IC_A); SLC26A4 and CLCNKB for CD_IC_B; GZMA and GZMB for NK_cells; CD3D, CD3E, and CD3G for T_cells; CD14, S100A8, and S100A9 for Macrophage_1; CD14 and FCER1A for Macrophage_2; CD79A and CD79B for B_cells.

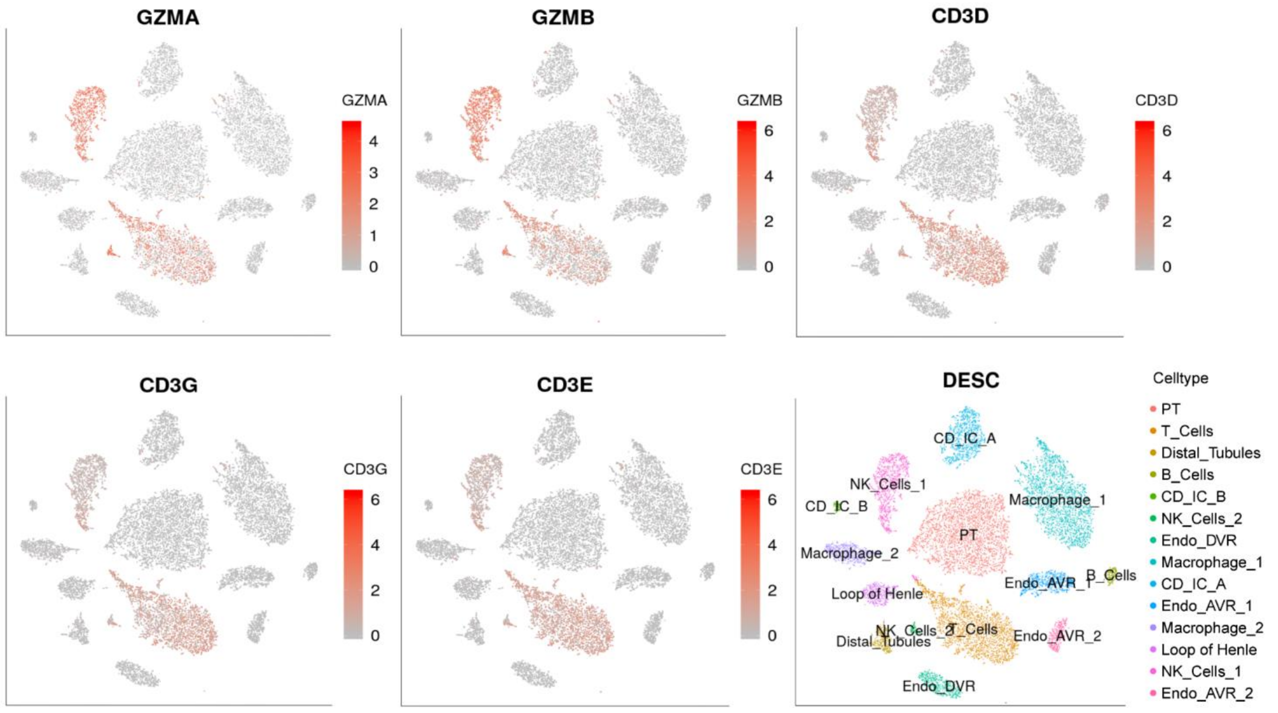


**Supplementary Fig 11.** Gene expression feature plot of the human kidney data for NK and T cell marker genes for DESC clustering. CD3D, CD3E and CD3G are marker genes for T cells, but GZMA and GZMB are marker genes for both NK cells and T cells.

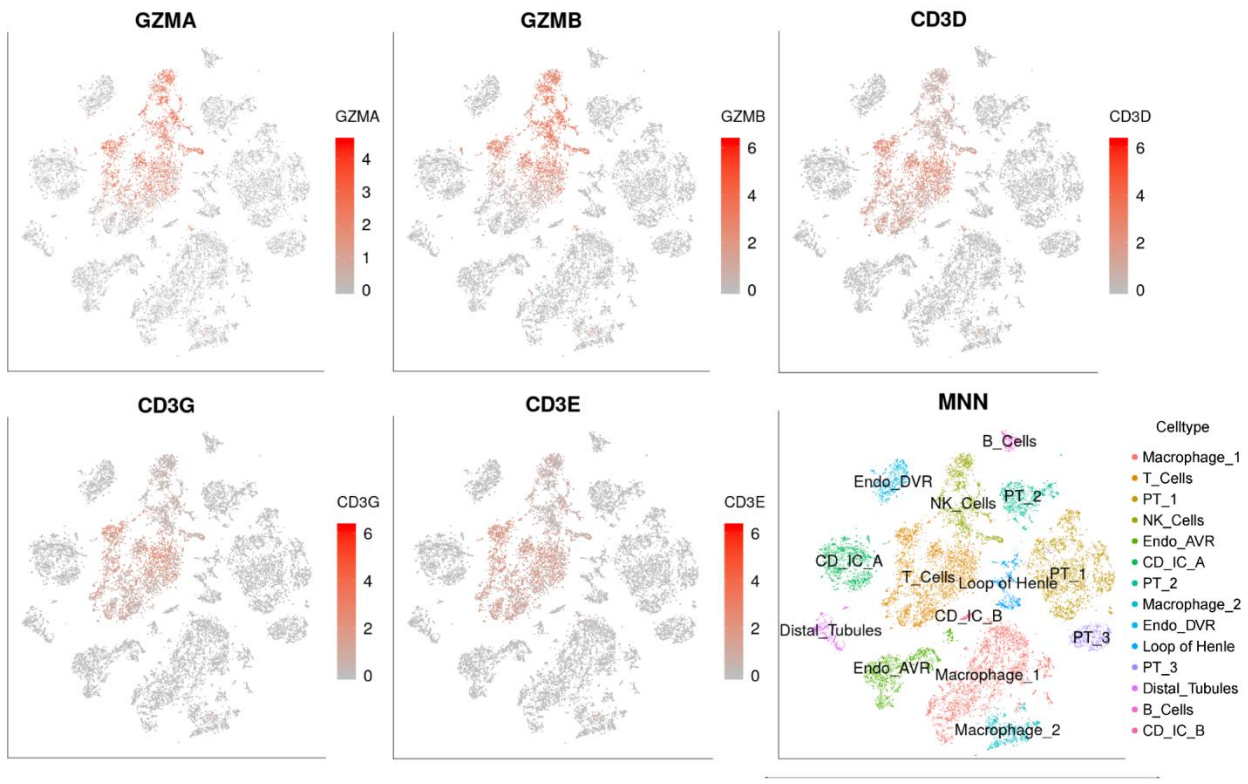


**Supplementary Fig 12.** Gene expression feature plot of the human kidney data for NK and T cell marker genes for MNN clustering. Note that CD3D, CD3E and CD3G are marker genes for T cells, but GZMA and GZMB are marker genes for both NK cells and T cells.

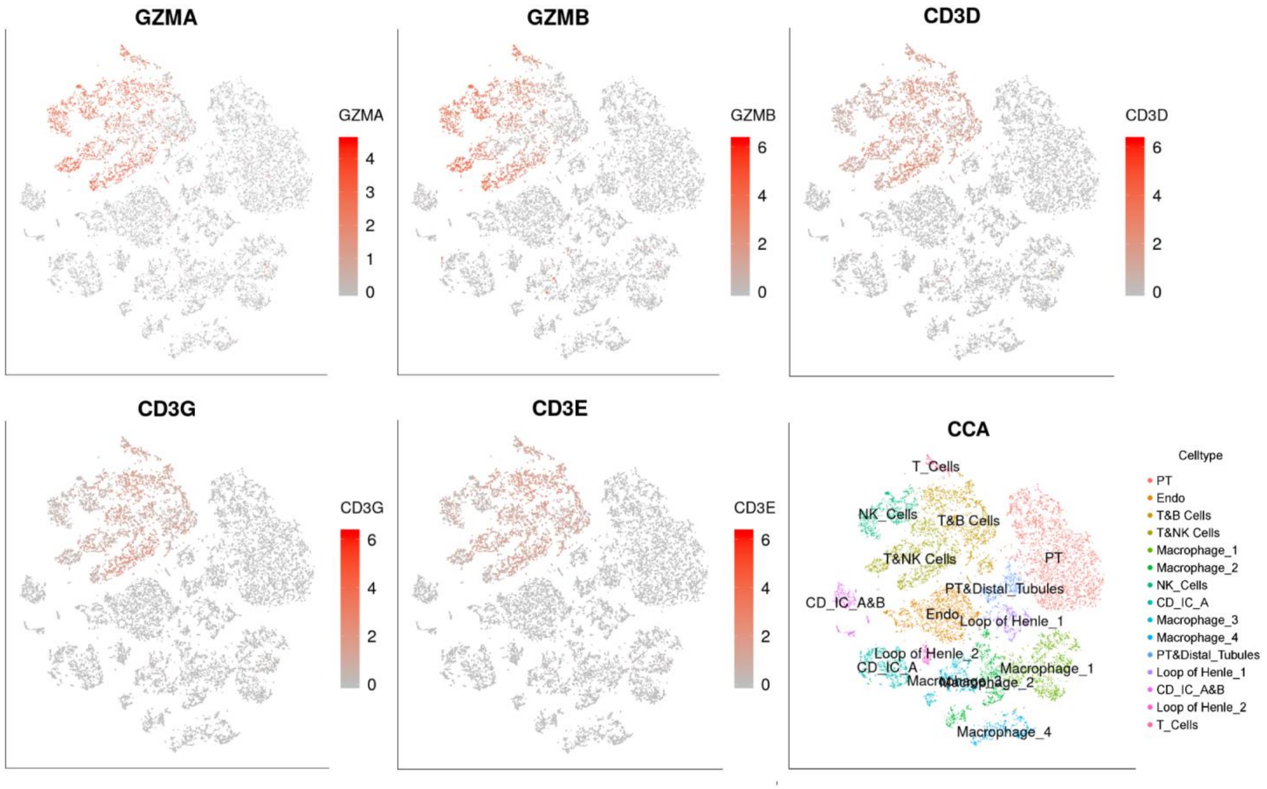


**Supplementary Fig 13.** Gene expression feature plot of the human kidney data for NK and T cell marker genes for CCA clustering. Note that CD3D, CD3E and CD3G are marker genes for T cells, but GZMA and GZMB are marker genes for both NK cells and T cells.

**Supplementary Note 5: analysis of the human PBMC data**

This dataset was generated by Kang et al. (2018) Multiplexed droplet single-cell RNA-sequencing using natural genetic variation. Nature Biotechnology 36(1):89-94.

The data were downloaded from GEO (https://ftp.ncbi.nlm.nih.gov/geo/series/GSE96nnn/GSE96583/suppl/), which include the raw gene count matrix, meta.data (t-SNE coordinates, ClusterID, celltype, and BatchID etc.) reported in the original paper. The downloaded data include 29,065 cells and 35,636 genes.

Cell filtering criteria: 1) eliminated cells that were labeled as multiplets and doublet.

Gene filtering criteria: 1) a gene was eliminated if the number of cells expressing this gene is <10.

Data processing: 1) gene expression levels for each cell was normalized using the "scanpy.api.normalize_per_cell" function in scanpy with counts_per_cell_after =10,000; 2) top 1,000 highly variable genes were selected using the "scanpy.api.pp.filter_genes_dispersion" function in scanpy; 3) normalized gene expression for the selected top 1,000 highly variable genes was then transformed using log(1+x) transformation with natural logarithm; 4) the expression is further standardized to a z-score, and the standardized gene expression values were used as input for DESC.

After the above filtering and data processing, there were 24,679 cells ×1,000 highly variable genes remained in DESC analysis.

DESC analysis: we used two hidden layers with 128 nodes in the first hidden layer, and 32 nodes in the second hidden layer. Other parameters were set default values. The final model was 1000-128-32-128-1000.

Other analyses: For CCA, we used the same number of cells, and performed analysis following Seurat's CCA tutorial (https://satijalab.org/seurat/immune_alignment.html). Specifically, we selected top 1,000 highly variable genes for each condition (control, stimulus), pooled these genes together and then removed genes that are expressed only in one condition. For MNN, we used the same number of cells, and highly variable genes were selected using function `scanpy.api.pp.fiter_gene_ dispersion` in python module scanpy with default parameters.

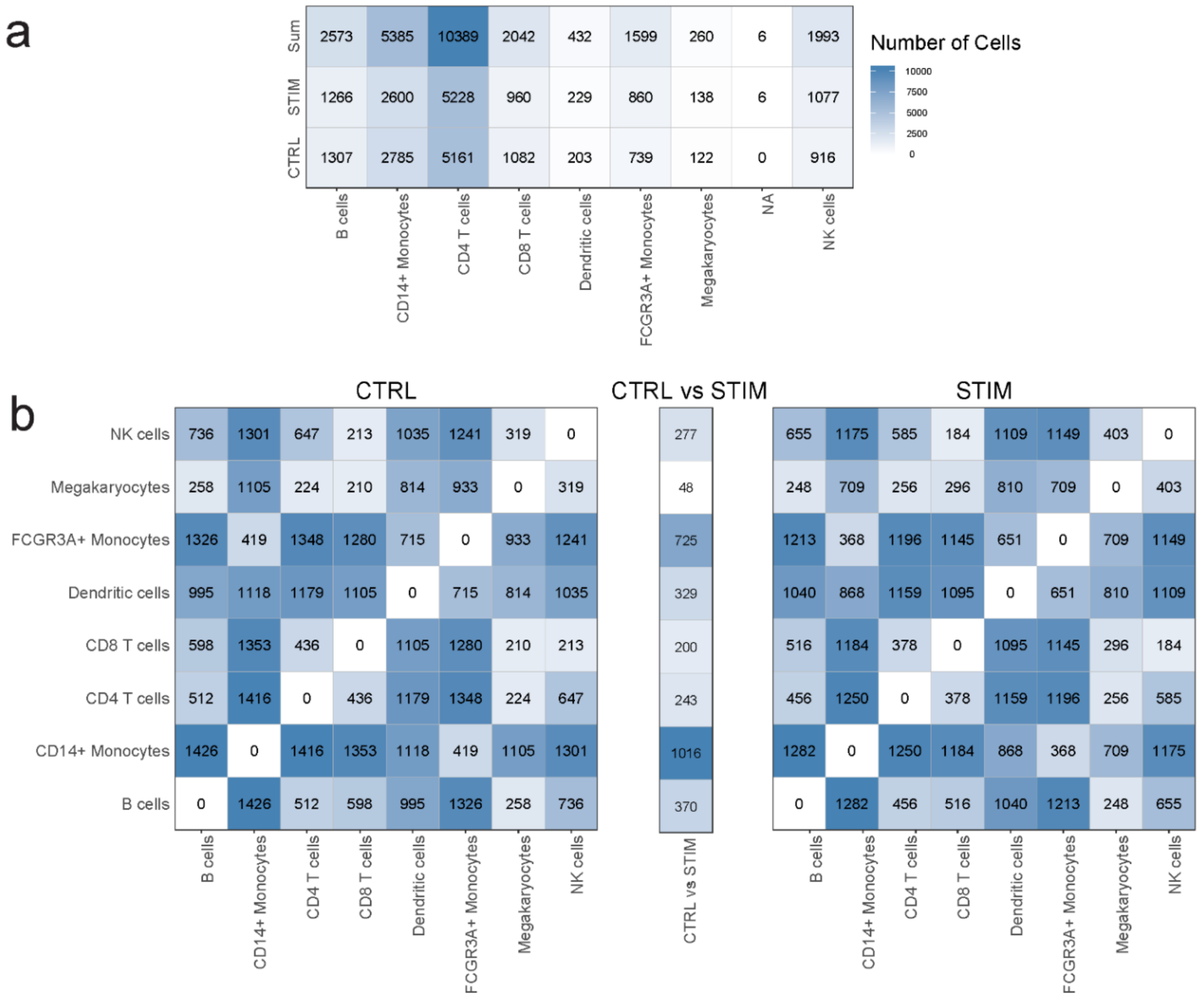


**Supplementary Fig 14. (a)** Number of cells in each cell type. Cell types were based on information provided in the original paper (Kang et al. 2018). **(b)** Number of differentially expressed genes (FDR adjusted p-value <0.01) between different cell types in the control group (left), the stimulus group (right), and differentially expressed genes between the control and the stimulus group within the same cell type (middle).

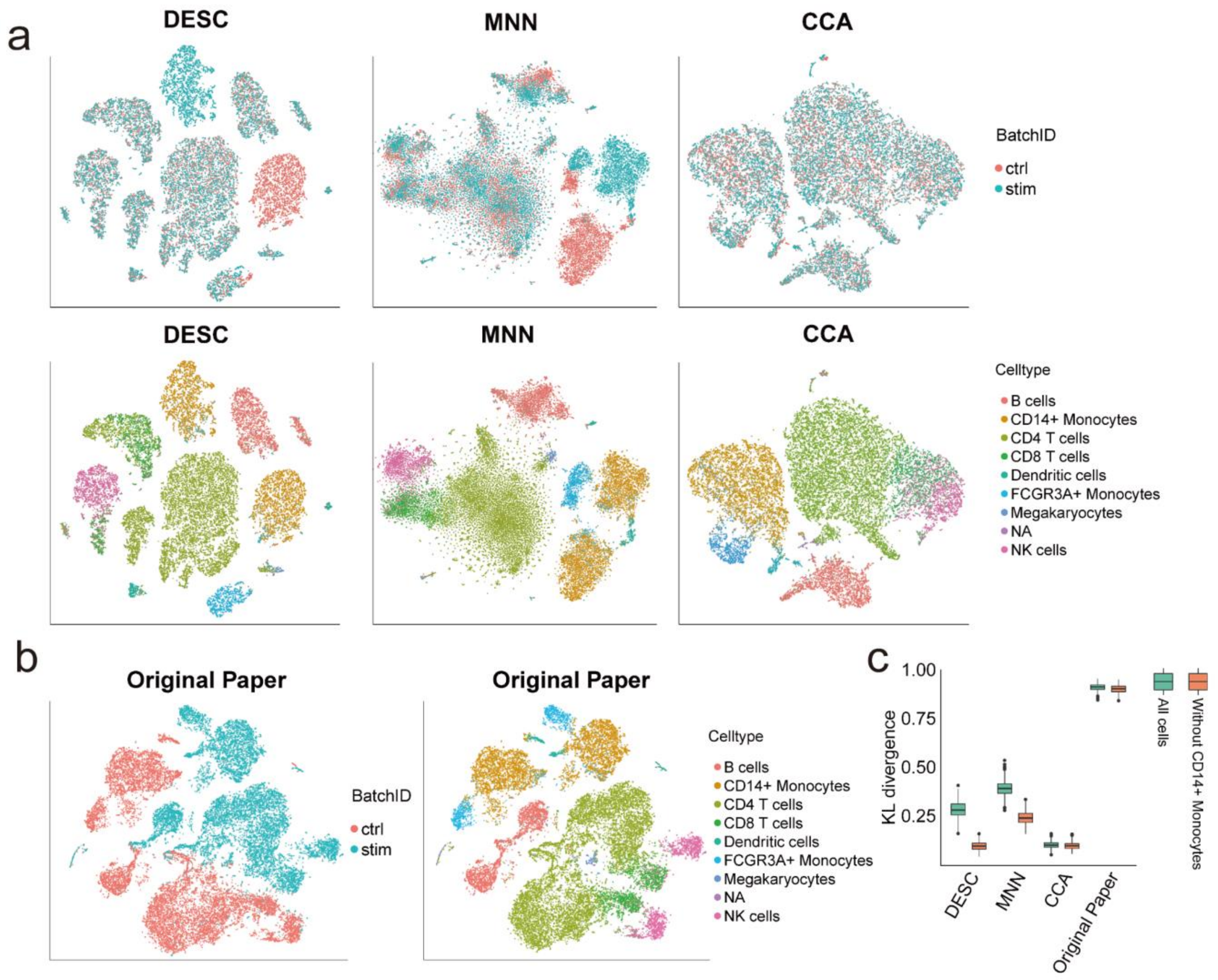


**Supplementary Fig 15.** Comparison of KL-divergence of DESC, MNN, CCA, and original analysis result on the PBMC data. **(a)** The t-SNE plots of each method, where the cells were colored by BatchID and cell type. **(b)** The t-SNE coordinates were from the original paper, and the cells were colored by BatchID (left) and cell type (right). The cell type label was determined by the original paper. **(c)** The KL-divergence calculated using all cells (colored by green), and the KL-divergence calculated using non CD14+ Monocytes (colored by red). The decreased KL divergence of DESC when CD14+ Monocytes were eliminated indicates that technical batch effect was effectively removed in the absence of CD14+ Monocytes. The KL divergence of MNN is larger than DESC when CD14+ Monocytes were eliminated, indicating that it might be less effective in removing technical batch effect than DESC. CCA has similar KL divergence irrespective of CD14+ Monocytes, indicating that it may have overcorrected batch effect, leading to the loss of biological variations between cells. The original analysis had substantially larger KL divergence than DESC, MNN, and CCA, indicating strong batch effect.

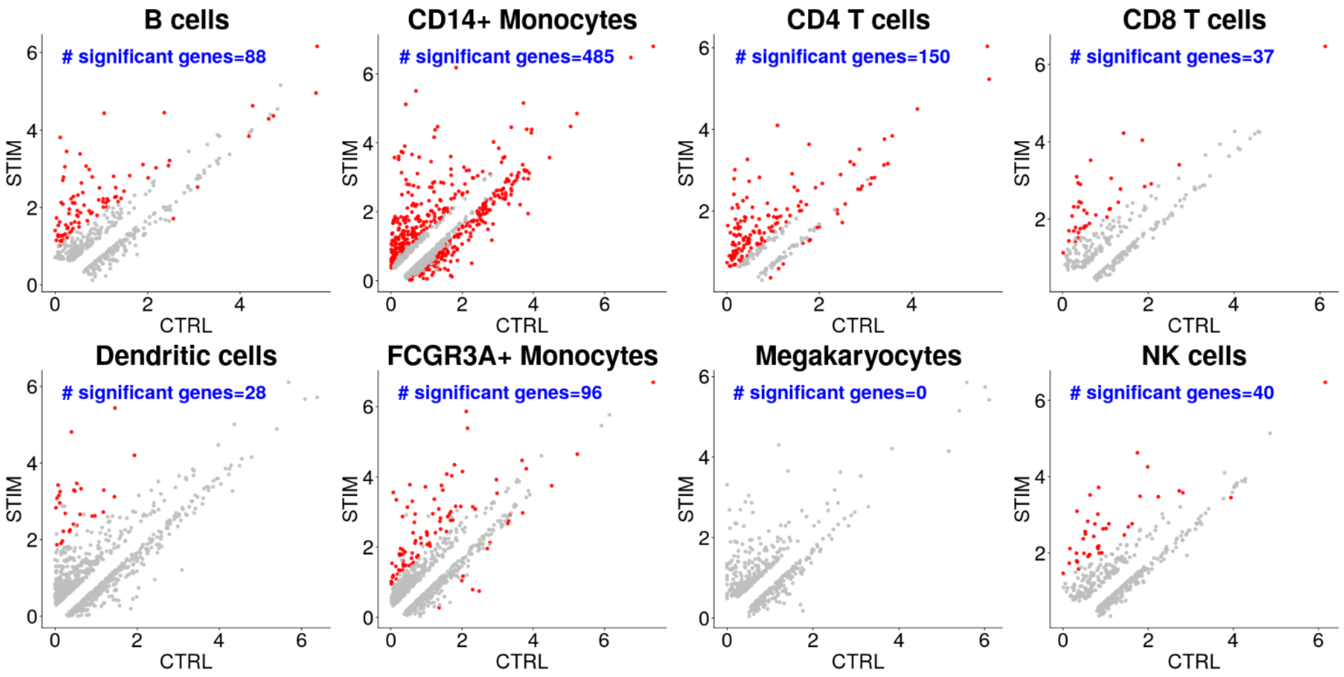


**Supplementary Fig 16.** Comparison of gene expression levels between control and stimulus conditions on the PBCM data. Displayed are the average gene expression across all cells in each condition for each cell type. Highlighted are differentially expressed genes with FDR adjusted p-value<$10^{-50}$.

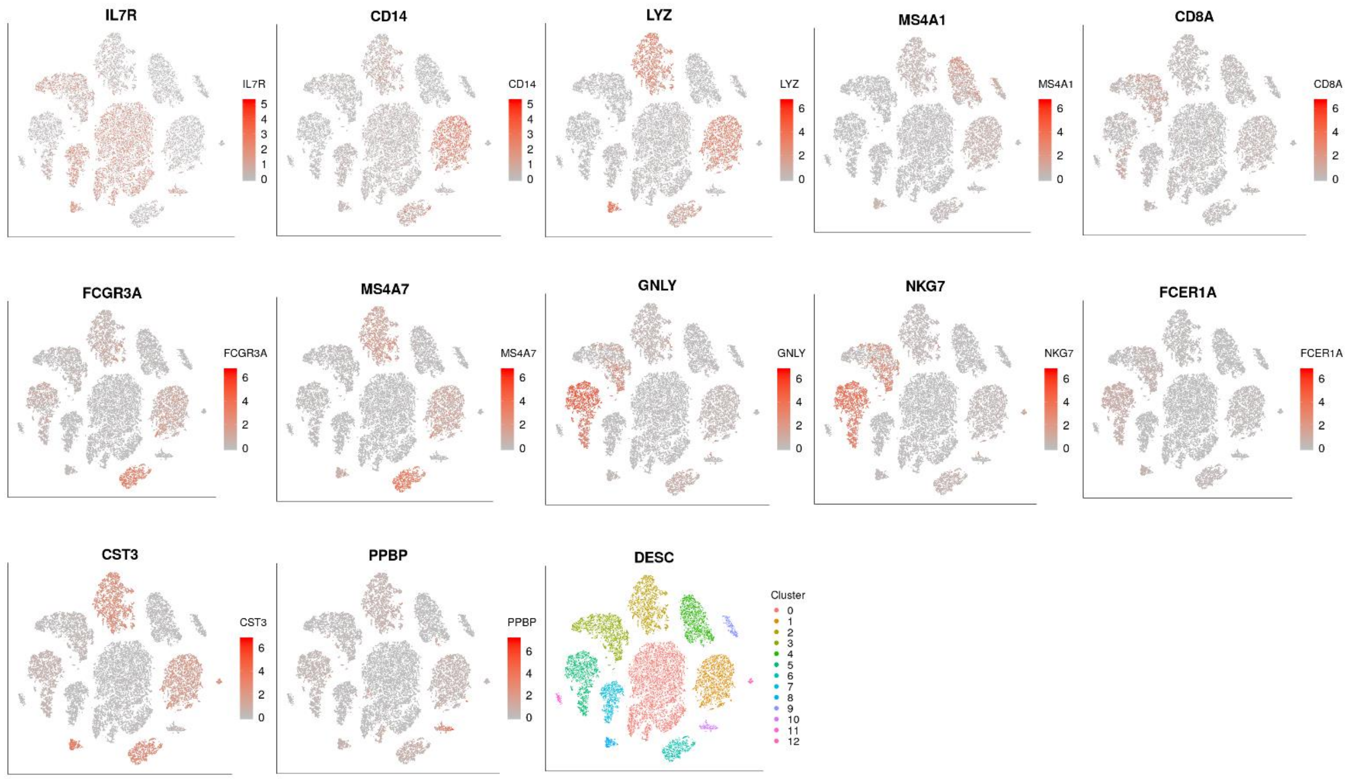


**Supplementary Fig 17.** Gene expression feature plots for cell-type specific marker genes for DESC clustering (resolution =0.8) on the PBMC data. IL7R (CD4 T cells marker), CD14 (CD14+ Monocyte marker), LYZ (CD14+ Monocyte marker), MS4A1 (B cells marker), CD8A (CD8 T cell marker), FCGR3A (FCGR3A+ monocyte marker), MS4A7 (FCGR3A+ Monocytes marker), GNLY (NK cells marker), NKG7 (NK cell marker), FCER1A (Dendritic Cells marker), CST3 (Dendritic Cells marker), PPBP (Megakaryocytes marker).

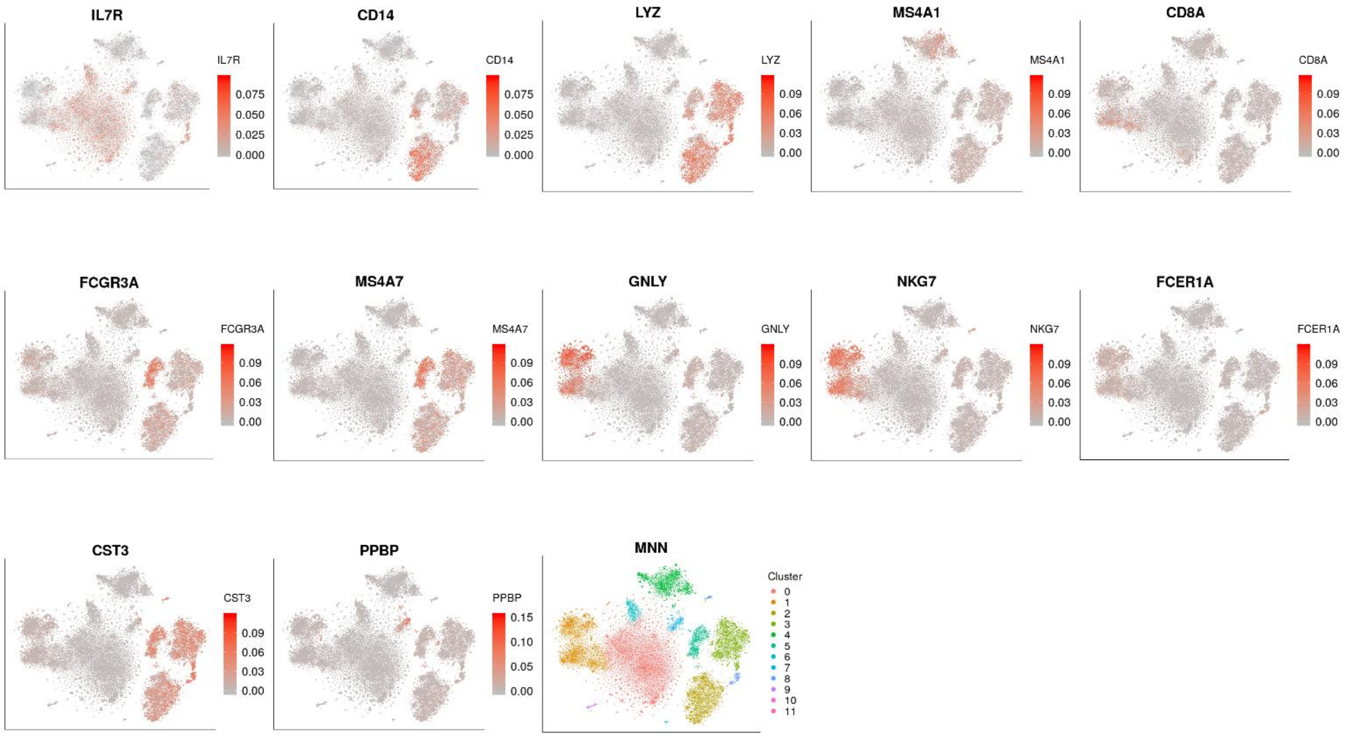


**Supplementary Fig 18.** Gene expression feature plots for cell-type specific marker genes for MNN clustering (resolution =0.4) on PBMC data. IL7R (CD4 T cells marker), CD14 (CD14+ Monocyte marker), LYZ (CD14+ Monocyte marker), MS4A1 (B cells marker), CD8A (CD8 T cell marker), FCGR3A (FCGR3A+ monocyte marker), MS4A7 (FCGR3A+ Monocytes marker), GNLY (NK cells marker), NKG7 (NK cell marker), FCER1A (Dendritic Cells marker), CST3 (Dendritic Cells marker), PPBP (Megakaryocytes marker).

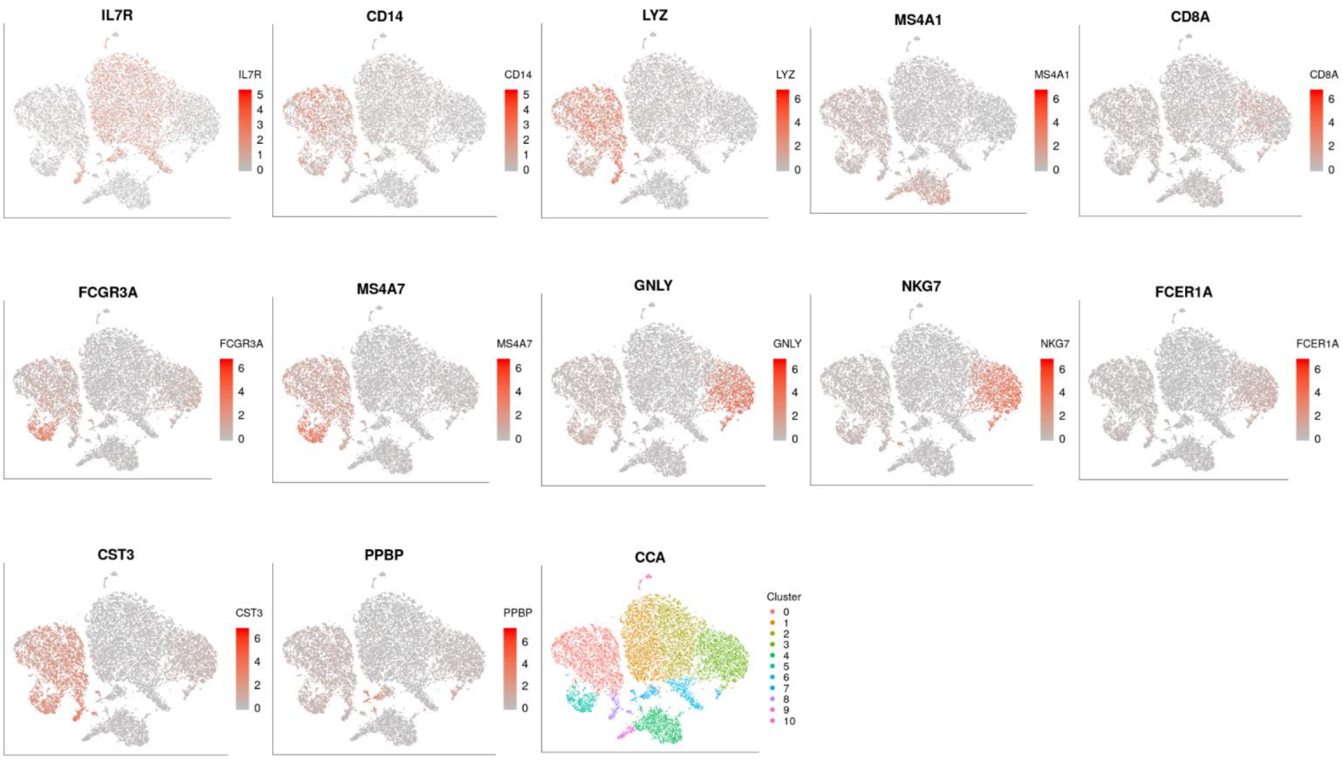


**Supplementary Fig 19.** Gene expression feature plots for cell-type specific marker genes for CCA clustering (resolution =0.4) on the PBMC data. IL7R (CD4 T cells marker), CD14 (CD14+ Monocyte marker), LYZ (CD14+ Monocyte marker), MS4A1 (B cells marker), CD8A (CD8 T cell marker), FCGR3A (FCGR3A+ monocyte marker), MS4A7 (FCGR3A+ Monocytes marker), GNLY (NK cells marker), NKG7 (NK cell marker), FCER1A (Dendritic Cells marker), CST3 (Dendritic Cells marker), PPBP (Megakaryocytes marker).

**Supplementary Note 6: analysis of the data with 1.3 million cells.**

The data were download from 10X website (https://support.10xgenomics.com/single-cell-gene-expression/datasets/1.3.0/1M_neurons). The original data include 1,306,127 cells and 27,998 genes.

Cell filtering criteria: 1) eliminated cells with gene counts <200;

Gene filtering criteria: 1) a gene was eliminated if the number of cells expressing this gene is <20.

Data processing: 1) gene expression levels for each cell was normalized using the "scanpy.api.normalize_per_cell" function in scanpy with counts_per_cell_after =10,000; 2) top 1,000 highly variable genes were selected using the "scanpy.api.pp.filter_genes_dispersion" function in scanpy; 3) normalized gene expression for the selected top 1,000 highly variable genes was then transformed using log(1+x) transformation with natural logarithm; 4) the expression is further standardized to a z-score, and the standardized gene expression values were used as input for DESC. After the above filtering and data processing, there are 1,292,537cells ×1000 highly variable genes remained in DESC.

DESC analysis: we used two hidden layers with 64 nodes in the first hidden layer, and 32 nodes in the second hidden layer. The parameters we used as n_neightbors=15, batch_size=20000, tol=0.008, louvain_resolution=0.2, use_GPU=True, is_stacked=False, pretrain_epochs=10, epochs_fit=2. Other parameters were set at default values. The final model is 1000-64-32-64-1000.

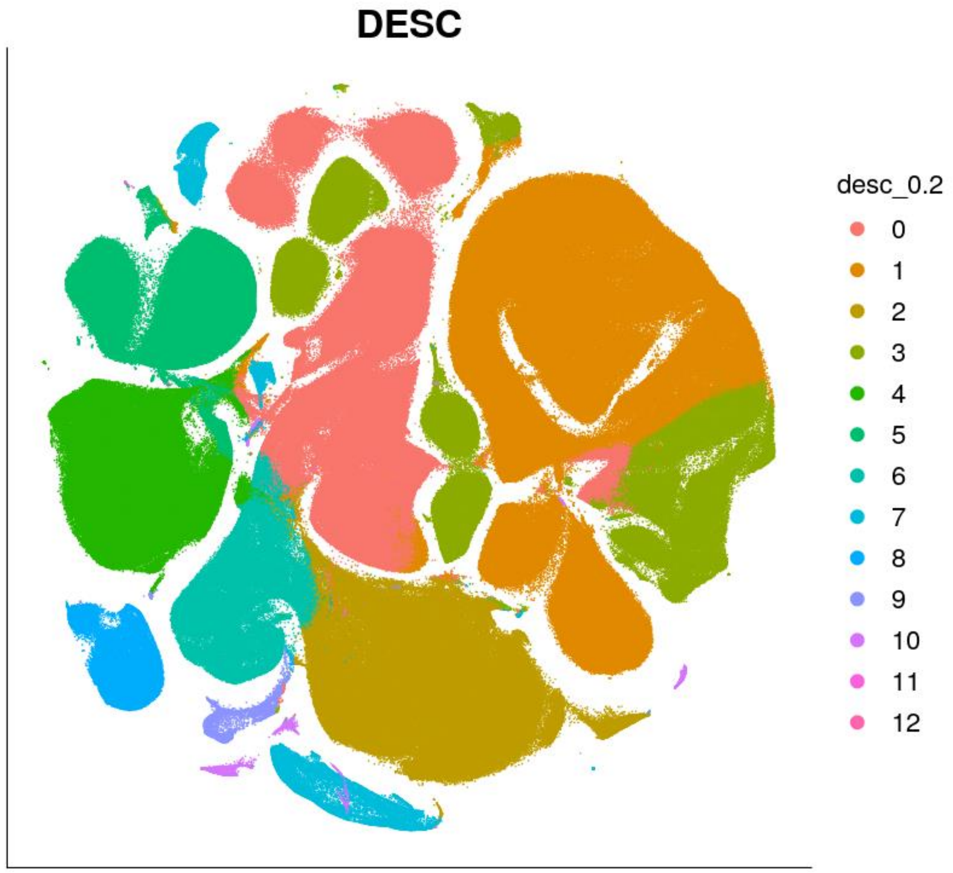


**Supplementary Fig 20.** Clustering result of DESC on the 1.3 million mouse data.

**Supplementary Note 7. Computing time and memory usage**

The data were download from 10X website (https://support.10xgenomics.com/single-cell-gene-expression/datasets/1.3.0/1M_neurons). The original data include 1,306,127 cells and 27,998 genes. For the sake of aesthetics, we abbreviate the number all cells as 1,300,000.

In order to compare computing time and memory usage for different methods and different numbers of cells, we randomly selected 1,000, 2,000, 5,000,10,000, 20,000, 40,000, 50,000, 100,000, 300,000, 500,000, 1,000,000, 1,300,000 cells from the above 1.3 million cell dataset.

We put **Fig 3e** here again for easy comparison of different methods.

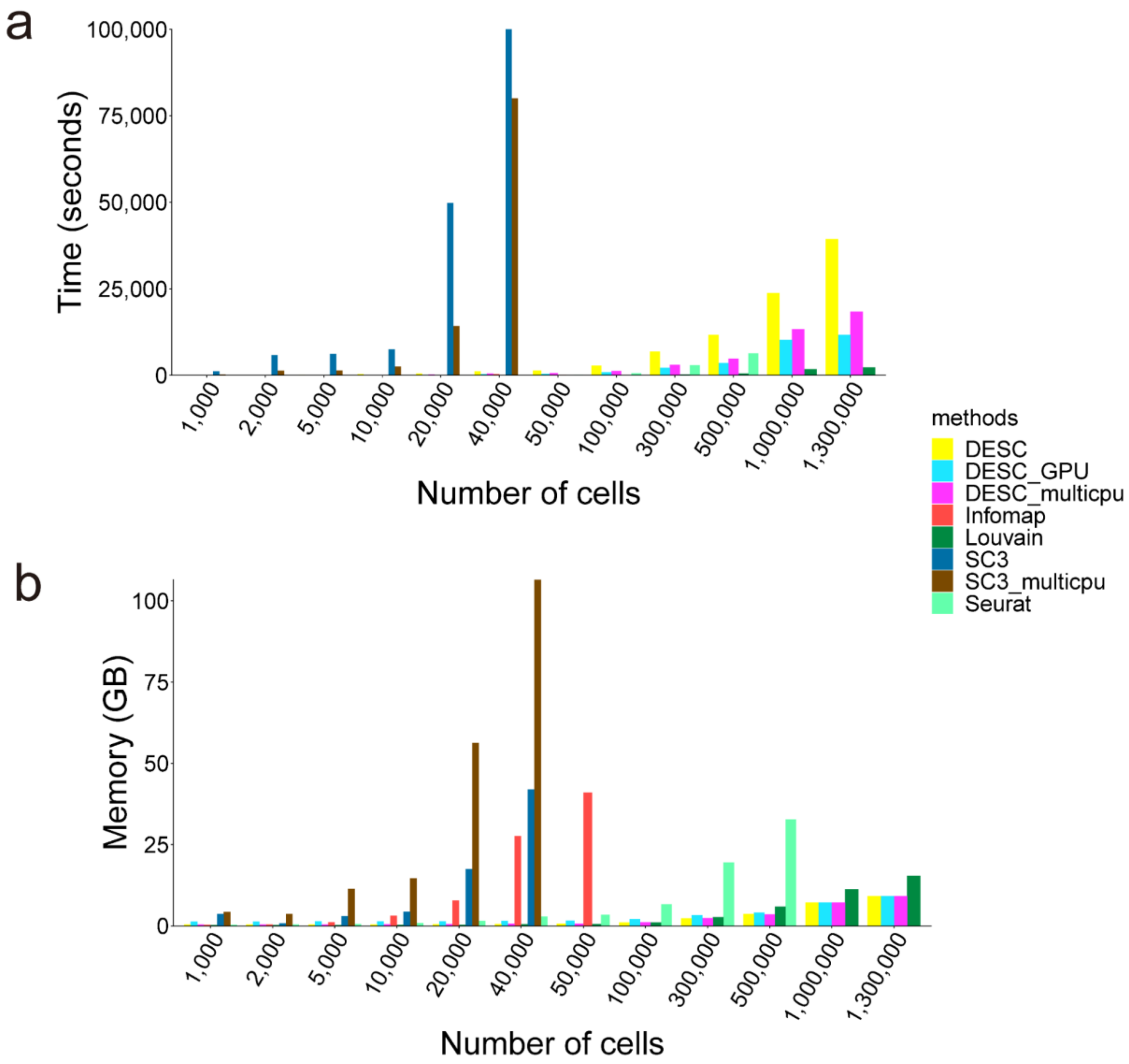


**Supplementary Fig 21**. Comparison of running time **(a)** and memory usage **(b)** of each method for datasets with various numbers of cells, where the cells were randomly sampled from the 1.3 million mouse brain dataset. DESC: used a single CPU; DESC_GPU: used GPU; DESC_multicpu: used 10 CPUs; SC3: used a single CPU; SC3_multicpu: used 5 CPUs. All other methods used a single CPU.

For each method, we monitored its memory use every second when the method was running, so we knew the memory use until the program ended. The memory we reported is the maximum memory use.

For DESC, DESC_GPU, DESC_multicpu and Louvain's method, we successfully completed analyses for all datasets. But for other methods, due to their memory issues, we only finished analysis for datasets with ≤500,000 cells (for Infomap and Seurat) or ≤400,000 (for SC3 and SC3_multicpu). There was a memory issue for SC3_multicpu when the number of cells is 40,000, so the runtime of 80,000 seconds and memory use of 100 GB were estimated values based on our experience of running SC3_multicpu for datasets with slightly smaller number of cells.

Although the running time for DESC with a single CPU is slow, we can take advantage of GPU's efficiency and speed up computing remarkably. More importantly, the memory use of DESC increases linearly with the increasing number of cells (**Supplementary Fig 23b**), which makes it a practical choice for large datasets.

All data analyses reported in this paper, except for the 1.3 million cells mouse brain data, were conducted on Ubuntu 18.04.1 LTS with 12 Intel® Core(TM) i7-8700K CPU @ 3.70GHz and 64GB memory. For the 1.3 million cells mouse brain dataset, we analyzed on Ubuntu 16.04.4 LTS with 32 Intel(R) Xeon(R) CPU E5-2620 v4 @ 2.10GHz and 128G memory.
